# CoRE-ATAC: A deep learning model for the functional classification of regulatory elements from single cell and bulk ATAC-seq data

**DOI:** 10.1101/2020.06.22.165183

**Authors:** Asa Thibodeau, Shubham Khetan, Alper Eroglu, Ryan Tewhey, Michael L. Stitzel, Duygu Ucar

**Affiliations:** The Jackson Laboratory for Genomic Medicine, Farmington, Connecticut, United States of America; Department of Genetics and Genome Sciences, University of Connecticut Health Center, Farmington, Connecticut, United States of America; The Jackson Laboratory, Bar Harbor, Maine, United States of America; Institute for Systems Genomics, University of Connecticut Health Center, Farmington, Connecticut, United States of America

**Keywords:** deep learning, ATAC-seq, snATAC-seq, *cis*-REs, enhancers, insulators

## Abstract

*Cis*-Regulatory elements (cis-REs) include promoters, enhancers, and insulators that regulate gene expression programs *via* binding of transcription factors. ATAC-seq technology effectively identifies active *cis*-REs in a given cell type (including from single cells) by mapping accessible chromatin at base-pair resolution. However, these maps are not immediately useful for inferring specific functions of *cis*-REs. For this purpose, we developed a deep learning framework (CoRE-ATAC) with novel data encoders that integrate DNA sequence (reference or personal genotypes) with ATAC-seq cut sites and read pileups. CoRE-ATAC was trained on 4 cell types (n=6 samples/replicates) and accurately predicted known *cis*-RE functions from 7 cell types (n=40 samples) that were not used in model training (mean average precision=0.80). CoRE-ATAC enhancer predictions from 19 human islet samples coincided with genetically modulated gain/loss of enhancer activity, which was confirmed by massively parallel reporter assays (MPRAs). Finally, CoRE-ATAC effectively inferred *cis*-RE function from aggregate single nucleus ATAC-seq (snATAC) data from human blood-derived immune cells that overlapped with known functional annotations in sorted immune cells, which established the efficacy of these models to study cis-RE functions of rare cells without the need for cell sorting. ATAC-seq maps from primary human cells reveal individual- and cell-specific variation in *cis*-RE activity. CoRE-ATAC increases the functional resolution of these maps, a critical step for studying regulatory disruptions behind diseases.

**Author Summary:** Non-coding DNA sequences serve different functional roles to regulate gene expression. For these sequences to be active, they must be accessible for proteins and other factors to bind in order to carry out a specific regulatory function. Even so, mutations within these sequences or other regulatory events may modulate their activity or regulatory function. It is therefore critical that we identify these non-coding sequences and their specific regulatory function to fully understand how specific genes are regulated. Current sequencing technologies allow us to identify accessible sequences via chromatin accessibility maps from low cell numbers, enabling the study of clinical samples. However, determining the functional role associated with these sequences remains a challenge. Towards this goal, we harnessed the power of deep learning to unravel the intricacies of chromatin accessibility maps to infer their associated gene regulatory functions. We demonstrate that our method, CoRE-ATAC, can infer regulatory functions in diverse cell types, captures activity differences modulated by genetic mutations, and can be applied to accessibility maps of single cell clusters to infer regulatory functions of rare cell populations. These inferences will further our understanding of how genes are regulated and enable the study of these mechanisms as they relate to disease.

## Introduction

*Cis*-Regulatory Elements (*cis*-REs) are non-coding DNA sequences that can be bound by transcription factors (TFs) and can take on different functional roles (e.g., promoter, enhancer, or insulator) to regulate gene expression programs. Over 88% of disease associated single nucleotide polymorphisms (SNPs) from genome-wide association studies (GWAS) are within non-coding region of the genome[1]. In particular, these SNPs typically fall within cell-specific enhancer sequences and indirectly disrupt gene expression programs[2–4]. It is therefore critical to map *cis*-REs and their functions with increased precision to study how GWAS SNPs disrupt gene regulation in different cell types. Furthermore, genetic variation can impact the activity level of *cis*-REs[5–8]. Uncovering the functionality of such genetically-modulated *cis*-REs will help to guide *in-vivo* and *in-vitro* functional studies, by prioritizing genomic loci that are most likely to impact gene regulation for experimental validation. Furthermore, identifying *cis*-RE functions in clinical samples will help uncover individual-specific and disease-associated elements and their functional roles in pathogenesis.

ENCODE[9] and Roadmap[10] consortia successfully annotated *cis*-REs for 127 reference human cell/tissue types by profiling their epigenomes and analyzing them using a hidden markov model (HMM) based approach: ChromHMM[11]. ChromHMM integrates Chromatin Immunoprecipitation with sequencing (ChIP-seq) profiles of multiple histone modification marks and TFs to demarcate promoters, enhancers, insulators and other *cis*-RE functions. These reference epigenomes are valuable resources to uncover and study disease-relevant and cell-type-specific *cis*-REs. However, these maps serve as references and do not capture individual- or conditionspecific (e.g., activated cells) *cis*-REs. Moreover, these references do not include *cis*-REs of less frequently studied and/or rarer cells that are gaining attention with the advances in single cell profiling techniques. Although ChromHMM is very effective in inferring *cis*-RE function, it requires five or more ChIP-seq assays to be generated from the same sample, which is not always feasible due to the cost (both antibody and sequencing) and cell numbers required for ChIP-seq assays (10^4^-10^6^ cells per experiment).

An alternative strategy for genome-wide interrogation of *cis*-REs is through chromatin accessibility profiling, which identifies open chromatin regions that are accessible for binding of TFs or other regulatory proteins/RNAs. One of the most recent and most frequently used methods for identifying open chromatin regions is Assay for Transposase Accessible Chromatin using Sequencing (ATAC-seq)[12, 13]. ATAC-seq utilizes Tn5 transposase to cleave accessible DNA into fragments, which are sequenced to generate a genome-wide map of open chromatin (Fig 1 Top). The low input material needed to generate ATAC-seq libraries allows it to be applied in clinical samples, making it an ideal assay for studying *cis*-REs in individual epigenomes in health and disease. Furthermore, recent developments have enabled the generation of high-quality ATAC-seq maps from single cells (i.e., single nucleus ATAC-seq (snATAC))[14] and generate epigenomic maps from clinical samples of limited quantity. Advances in snATAC technology enables researchers to study chromatin accessibility maps at unprecedented resolution by detecting rarer cell types and by studying epigenomes of cells without the need to sort them, which could affect gene expression programs. Despite these promises of snATAC-seq data, there are not many studies that show that machine learning models built from bulk ATAC-seq data can also be used for predictions on aggregated snATAC-seq data. ATAC-seq is widely used to infer *cis*-REs from diverse cell types and cell states [6, 12, 15–20], however, the functionality of these *cis*-REs (e.g., enhancers, promoters, insulators) cannot be inferred from these assays without the help of new computational methods that can fully interrogate ATAC-seq data features and integrate it with DNA sequence features.

**Fig 1.**
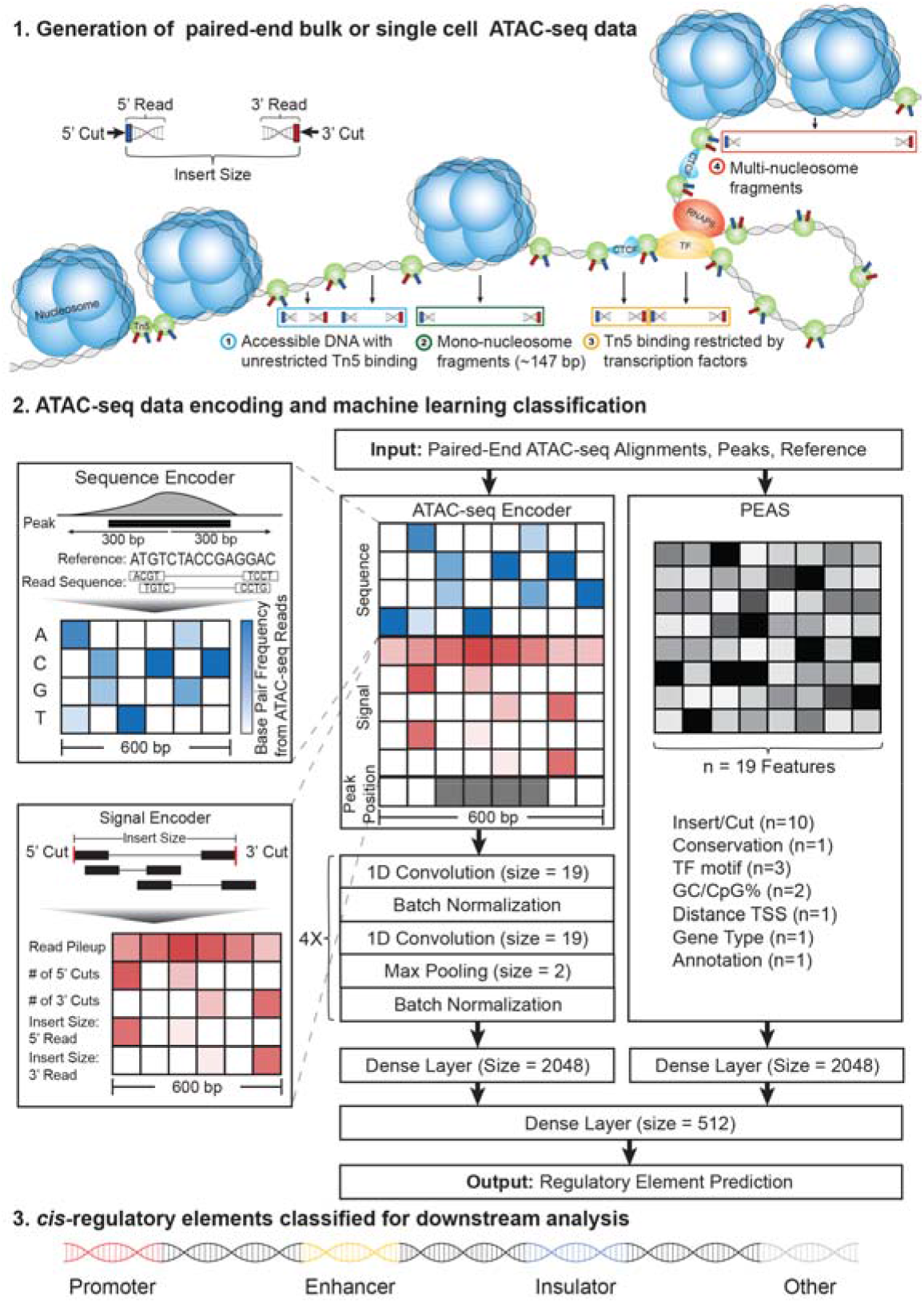
Overview of the CoRE-ATAC framework. Paired-end ATAC-seq data captures different cut and insert size distributions corresponding to the presence or absence of nucleosomes or TFs. ATAC-seq data is encoded into a 10×600 matrix and 19 data features from PEAS algorithm to predict the functionality of an open chromatin region, using both novel and manually selected features. In the final step, CoRE-ATAC classifies REs into 4 functional classes: promoter, enhancer, insulator, and other.

Previously we developed a method for predicting enhancers from bulk ATAC-seq data using a neural network (NN) model on extracted data features (n=24), namely PEAS[21]. Although PEAS was effective in predicting enhancers from ATAC-seq maps, it primarily utilized manually extracted features; potentially missing novel predictive features that could be hidden within the read count/cut site distributions of ATAC-seq data. Furthermore, PEAS was primarily developed to predict enhancer sequences, leaving the open question of whether other types of *cis*-REs can be predicted from ATAC-seq maps. To overcome these limitations, we turned to deep learning. Deep learning methods have revolutionized the machine learning field due to their ability to learn novel data features and have been successfully applied in genomics to predict chromatin accessibility[22] and enhancers[23, 24] from DNA sequence. We developed a deep learning framework, namely Classification of Regulatory Elements with ATAC-seq (CoRE-ATAC) (Fig 1 middle and bottom) to harness the power of deep learning for inferring the regulatory function of open chromatin regions. Core-ATAC integrates DNA sequence data with chromatin accessibility data (single cell or bulk) using a novel ATAC-seq data encoder that is designed to be able to integrate an individual’s genotype to personalize *cis*-RE predictions, especially for loci with genetically modulated regulatory activity (e.g., eQTLs, caQTLs).

In this study, we present CoRE-ATAC and evaluate its ability to predict *cis*-RE function (promoters, enhancers, insulators, and others) in 11 different cell types across 46 bulk ATAC-seq samples/replicates (Table 1) and demonstrate that CoRE-ATAC is a robust method that consistently predicts *cis*-REs with high average precision (mean micro average precision = 0.8) irrespective of whether the cell type was used in model training. We observed that CoRE-ATAC predictions recapitulated ChromHMM enhancers from the cognate cell type as well as *cis*-REs inferred *via* different assays (CAGE[25] and STARR-seq[26]). We compared CoRE-ATAC *cis*-RE predictions in human islet samples to *cis*-RE activity from massively parallel reporter assays (MPRA)[27] to demonstrate that CoRE-ATAC can predict the loss/gain of *cis*-RE activity linked to genetic variation. Finally, we showed that models built from bulk ATAC-seq data are also predictive on cell clusters from snATAC-seq data, by predicting *cis*-REs from snATAC-seq data in human Peripheral Blood Mononuclear Cells[14] for 7 blood-derived immune cell type clusters. Enhancers inferred from PBMC snATAC-seq data captured the majority of super enhancers in these immune cell subsets[2, 4] (i.e., cell-type-specific enhancers) and were enriched in SNPs for diseases of the cognate cell/tissue type. These analyses demonstrate the potential of CoRE-ATAC to annotate open chromatin regions from both bulk ATAC-seq and cell clusters from snATAC-seq samples, which will ultimately improve our understanding of how gene expression programs are regulated at the individual and cell-specific level and how this regulation is disrupted in pathologies.

**Table 1.** ATAC-seq samples used in model training and evaluation.

| Cell Type | Usage | # | Data Accession Id |
| --- | --- | --- | --- |
| CD14<sup>+</sup> | Model Training | 2 | EGAS00001002605 |
| K562 | Model Training | 2 | GSE121993 |
| HSMM | Model Training | 1 | GSE109828 |
| GM12878 | Model Training | 1 | GSE47753 |
| Pancreatic Islets | Cross-cell Predictions | 19 | SRP117935 |
| Naïve CD8<sup>+</sup> T | Cross-cell Predictions | 6 | EGAS00001002605 |
| PBMC | Cross-cell Predictions | 6 | EGAS00001002605 |
| Naïve CD8<sup>+</sup> T | Cross-cell Predictions | 4 | GSE118189 |
| MCF7 | Cross-cell Predictions | 2 | GSE97583 |
| A549 | Cross-cell Predictions | 1 | GSE117089 |
| CD4<sup>+</sup> T | Cross-cell Predictions | 1 | GSE47753 |
| EndoC | Cross-cell Predictions | 1 | GSE118588 |
| PBMC | snATAC Predictions | 1 | GSE129785 |

## Results

### CoRE-ATAC predicts functional annotations for *cis*-REs

Annotation of ATAC-seq peaks using ChromHMM states revealed that between 16-52% are promoters, 18-54% are enhancers, 4-15% are insulators (for samples with insulator states in ChromHMM) and 5-55% are other functional annotations (Fig 2a). The functional diversity in these annotations establishes a need for functionally annotating ATAC-seq open chromatin maps. Furthermore ChromHMM states are effective as baseline references to assess predictive performances of functional annotations[21] compared to alternatives (e.g., CAGE[25] or P300 binding) since ChromHMM captures a wider array of functional states and detects a larger set of loci (e.g., enhancers) that can be used in model training. Previously we showed that ChromHMM captured the majority of both P300 and CAGE identified enhancers, whereas P300 and CAGE enhancers identified smaller but distinct subsets of enhancers[21]. Therefore, we decided to use ChromHMM annotations as the ground truth in our models (further discussed in Discussion).

**Fig 2.**
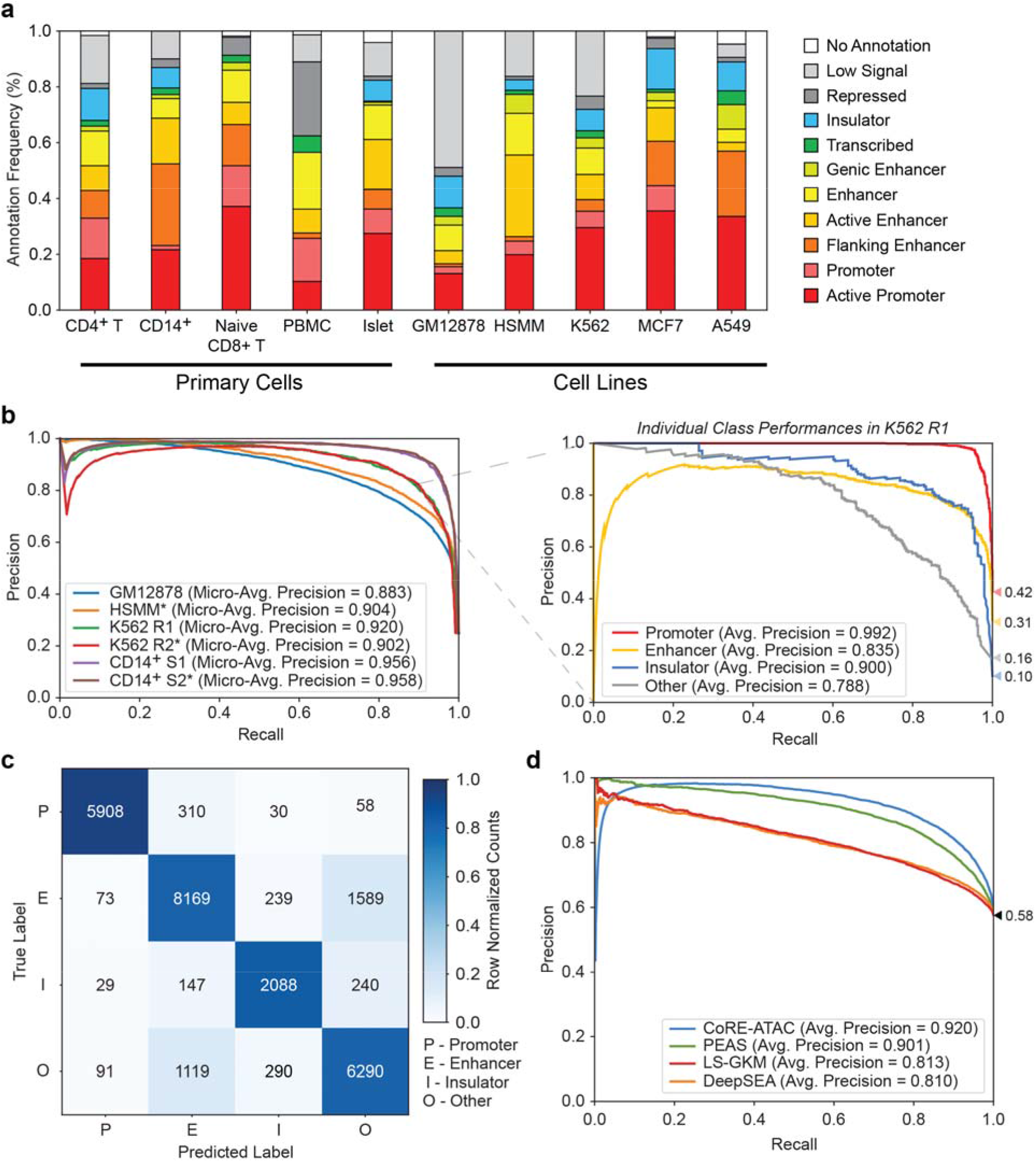
CoRE-ATAC outperforms sequence-based enhancer prediction methods. CoRE-ATAC predictions were evaluated using held out test data (chromosomes 3 and 11). (**a**) ChromHMM state distributions for different cell types used in this study. ATAC-seq open chromatin maps correspond to a multitude of *cis*-RE functional states, corresponding to promoters, enhancers, insulators, and other functional annotations. (**b**) Micro-average precision values (left) were calculated, summarizing the average precision values for individual class predictions for all cell types used in model training. A breakdown of individual class average precision scores is shown for K562 (right). (**c**) Combined confusion matrix of model predictions across all cell types used in model training. Note that models are predictive for all class labels. However, mispredictions were more frequently observed between enhancer and other functional classes. (**d**) Receiver operating characteristic (ROC) curves for different enhancer prediction models: CoRE-ATAC, PEAS, DeepSEA and LS-GKM. Note that CoRE-ATAC outperforms alternative methods.

Using ChromHMM annotations in cognate cell types as class labels, CoRE-ATAC was trained on GM12878, K562, HSMM, and CD14^+^ ATAC-seq samples and model performances were evaluated using held-out test data (i.e., regions within chromosomes 3 and 11) (Table 2). After identifying a concordant set of 10 functional states to use as a ground truth based on in-house and Roadmap ChromHMM states (S1 Fig and S2 Fig), we showed that among these, four states could be effectively discriminated from ATAC-seq maps (S3 Fig). We therefore annotated ATAC-seq peaks for three major functional *cis*-RE classes (promoters, enhancers, and insulators) using ChromHMM and combined other states into a fourth class, named “other” which captures states such as repressed, transcribed, and low signal/quiescent regions, covering ATAC-seq peaks that do not correspond to the other *cis*-REs (Methods) (S4 Fig). For the remainder of our analyses, we focus on models utilizing these four functional states.

**Table 2.**
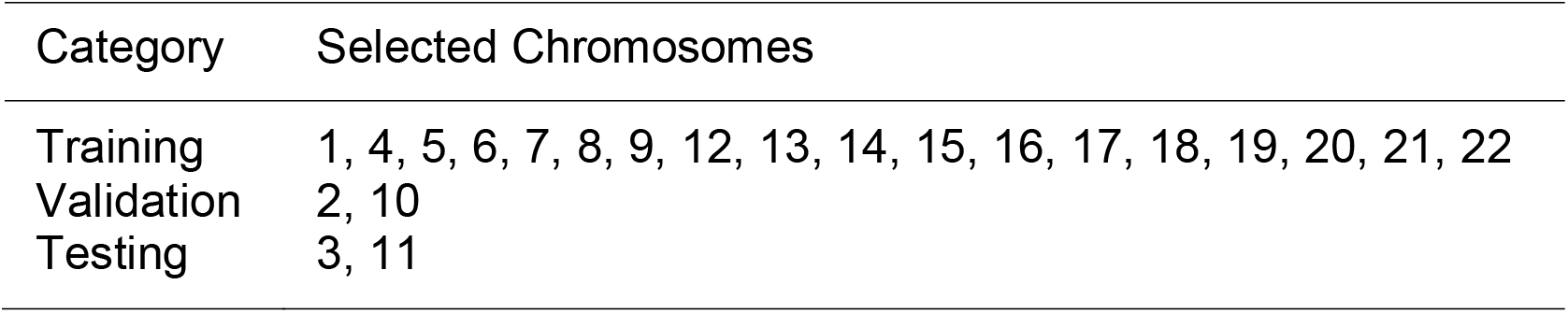
Chromosomes selected for training, validation, and test data.

CoRE-ATAC models evaluated on 4 cell types (Table 1) had high micro-average precision (0.88-0.96) across all samples with a combined accuracy of 84.20% (Fig 2b left and Fig 2c). Individual class performances also observed high average precision for all four classes of *cis*-REs (Fig 2b right, S5 Fig), establishing that high precision/accuracy values from CoRE-ATAC predictions are not driven by predictions of a single functional class. However, we noted that most mis-predictions were between enhancer and “other” classes (Fig 2c). Currently, the alternatives for inferring *cis*-RE function from ATAC-seq data are PEAS[21] and DNA sequence methods. We therefore compared the performance of CoRE-ATAC with two sequence based methods: DeepSEA[24] and LS-GKM[28], and our previous neural network based approach PEAS[21] using the same training and testing data for enhancer and “other” classes. CoRE-ATAC showed improvement over these alternative methods (Fig 2d, S6 Fig) (average precision=0.92 for held out test data), with PEAS following second in performance (average precision=0.90). Methods based solely on DNA sequence performed similarly to one another (S6 Fig) (average precision=0.81) but were not as effective as PEAS and CoRE-ATAC. The increased predictive performances of CoRE-ATAC and PEAS suggest that ATAC-seq signal (e.g., read/insert pileups) contains critical information for functionally classifying *cis*-REs that cannot be captured from DNA sequence alone. This analysis establishes that CoRE-ATAC improves upon DNA sequence-based approaches by capturing relevant and predictive features from the ATAC-seq signal.

### CoRE-ATAC predicts *cis*-RE functionality across cell types

We evaluated CoRE-ATAC’s ability to predict *cis*-RE function in 40 bulk ATAC-seq samples from 7 cell types (i.e., pancreatic islets, MCF7 breast cancer cell line, naïve CD8^+^ T, PBMC, CD4^+^ T, A549 and EndoC beta cell line) that were not used in model training (Table 1). CoRE-ATAC models were highly predictive for cell types that were not utilized in model training, showing ~0.80 micro-average precision across all samples (Fig 3a). The highest precision values were detected for promoters (~0.95), followed by enhancers (~0.76) (Fig 3a). Insulator annotations were only available in 4 of the 7 cell types tested (MCF7, A549, CD4^+^ T and islet), however, insulator states in islets were excluded from these analyses due to the poor quality of CTCF ChIP-seq data in islets (S7 Fig), which affected the performance assessment for islet samples. Among the remaining 3 cell types, known insulators (based on ChromHMM states) were predicted with high average precision (~0.78) (Fig 3a and S8 Fig for islets). As expected, the majority of insulator predictions resided within CTCF ChIP-seq peaks (S9 Fig). To further show the functional relevance of CoRE-ATAC insulators that were either not annotated by ChromHMM (i.e., cell types that are missing insulator states) or were misclassifications compared to ChromHMM insulators, we conducted de novo motif enrichment analysis using HOMER[29]. CTCF motif was the most enriched sequence for these loci with a very significant p-value (P-value < 1e-27180) (Fig 3b), suggesting that CoRE-ATAC predictions capture CTCF insulator sequences.

**Fig 3.**
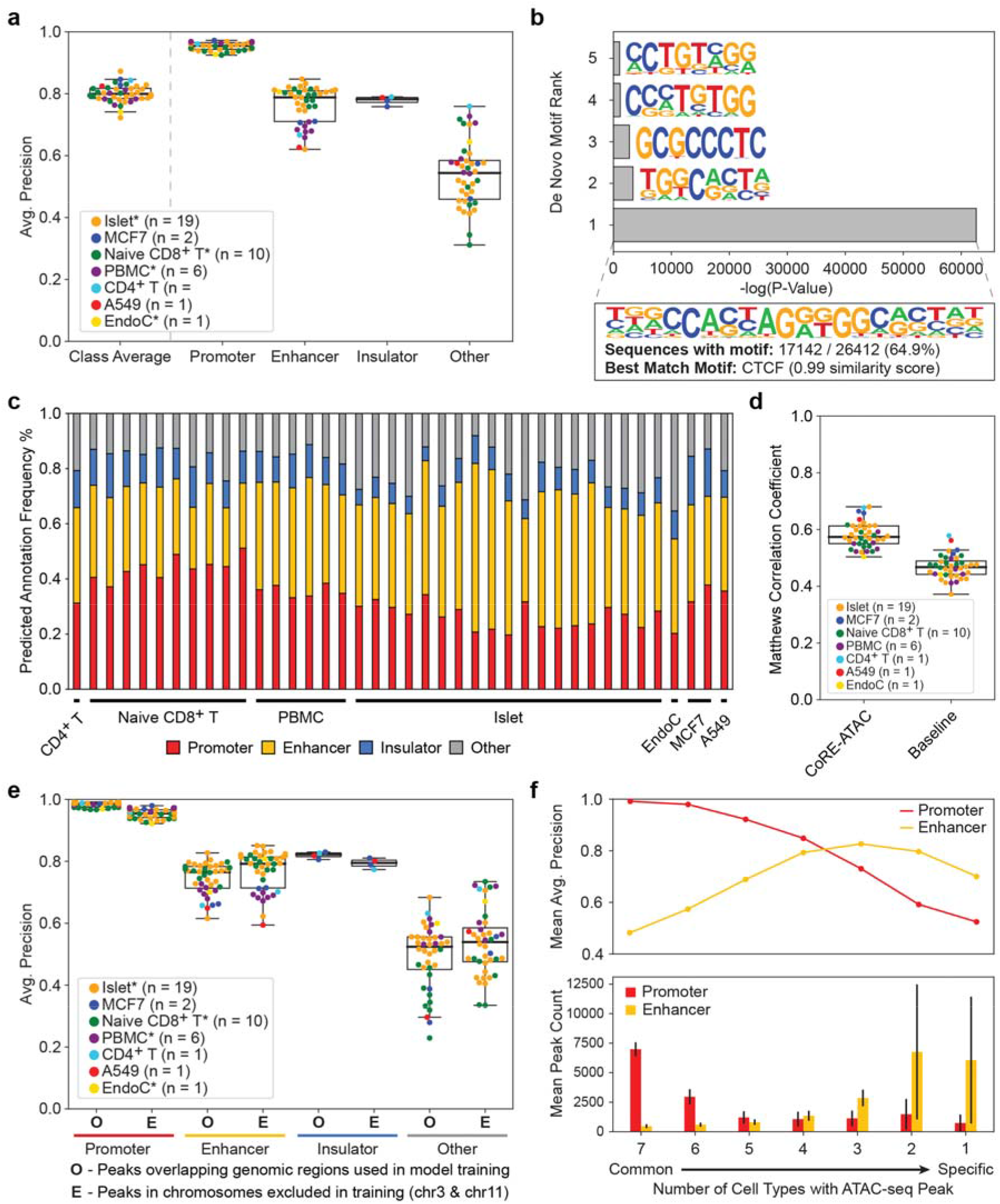
CoRE-ATAC can predict REs across cell-types. CoRE-ATAC was evaluated in 7 cell types using 40 samples that are not used in model training. (**a**) Average precision scores for predicting REs. Micro-average precision was used to calculate class average scores. Co-RE ATAC is predictive across cell types and different functional classes with an exception of insulators in islets, which is due to CTCF ChIP-seq quality in islets. (**b**) De novo motif enrichment results for regions predicted as insulators by CoRE-ATAC but were not annotated as insulators by ChromHMM. Note that these regions are significantly enriched for the CTCF motif (0.983 similarity), suggesting that CoRE-ATAC insulator predictions are functionally relevant. (**c**) Distribution of CoRE-ATAC predictions. Prediction distributions are similar to those observed by ChromHMM state annotations. (**d**) Comparison of CoRE-ATAC to baseline/naive predictions based on thresholds for distance to TSS, MACS2 FDR, and number of CTCF motifs. CoRE-ATAC improves upon baseline performances. (**e**) CoRE-ATAC performances for i) predictions overlapping regions used in model training (O), and ii) predictions within regions that are on held-out test chromosomes (E). Note the performance similarity between these two prediction categories across all classes. (**f**) CoRE-ATAC model performances (top) and the average number of promoters and enhancers observed (bottom) by cell-type-specificity. We observed that CoRE-ATAC was more effective in predicting common promoters and cell-type-specific enhancers, for which we had more examples represented in the data. CoRE-ATAC’s ability to predict cell-type-specific enhancers demonstrates its usefulness for interrogating individual and cell-type-specific enhancers.

As expected, functional annotation of ATAC-seq peaks from these 40 samples *via* CoRE-ATAC (Fig 3c) were similar to state distributions observed with ChromHMM (Fig 2a). In addition, we studied whether functional predictions with CoRE-ATAC can outperform a naïve annotation approach using distance to transcription start site (TSS), MACS2 FDR, and number of CTCF motifs (Methods). CoRE-ATAC annotations outperformed these naïve annotation approaches (Fig 3d, S10 Fig), observing ~0.58 Matthews correlation coefficient score for CoRE-ATAC compared to ~0.47 on the average across all samples. Matthews correlation coefficient was used to account for the fact that threshold-based approaches produce binary probabilities (i.e. 1.0 or 0.0) that can result in misleading average precision calculations.

A potential pitfall in cross-cell-type model evaluations is the use of the same genomic regions (e.g., same chromosome) in both model training and testing[30]. When the same region (not the same genomic data) is used in training and testing, a model might perform well simply because it “remembers” the specific DNA sequences used during training. To study whether our model suffers from this pitfall, we utilized the two chromosomes that were excluded from model training (chromosomes 3 and 11) and compared cross cell type predictions for these two chromosomes with regions that overlapped loci used in model training and observed comparable predictive performances (Fig 3e). These analyses suggest that CoRE-ATAC has learned a function from DNA sequence and chromatin accessibility signals that is transferable across genomic regions for predicting *cis*-RE functionality rather than learning to memorize specific DNA sequences.

Finally, we evaluated CoRE-ATAC’s cross-cell-type performance by stratifying ATAC-seq peaks by their cell-specificity across 7 cell types and compared prediction performances. CoRE-ATAC predicted common promoters with higher average precision than cell-type-specific promoters. Surprisingly, CoRE-ATAC predicted cell-type-specific enhancers more effectively compared to common enhancers (mean average precision 0.70 versus 0.48), emphasizing the utility of this method to study disease-relevant enhancers that are typically cell-type-specific[2] (Fig 3f top). The prediction bias observed for cell-specific promoters is likely due to the number of elements used in model training given that the majority of promoters are common among cell types (Fig 3f bottom), whereas the majority of enhancers are cell-type-specific.

### Core-ATAC can predict enhancers that are captured *via* different assays

To study whether CoRE-ATAC could detect enhancers identified by alternative methods, we compared ChromHMM annotations and CoRE-ATAC predictions in MCF7, A549, CD4^+^ T, and PBMC cell types to CAGE[25] enhancers identified by the FANTOM5 project[31, 32]. We observed that the majority of FANTOM enhancers overlapped with promoters and enhancers in both ChromHMM and CoRE-ATAC (Fig 4a). CoRE-ATAC enhancer predictions showed significant overlap with FANTOM enhancers (Fisher’s exact test p-values for all cell types < 1.26e-11). The similarity between ChromHMM and CoRE-ATAC further establishes that functional annotations *via* CoRE-ATAC using ATAC-seq data is a cost-effective alternative to annotations *via* ChromHMM that use multiple ChIP-seq assays.

**Fig 4.**
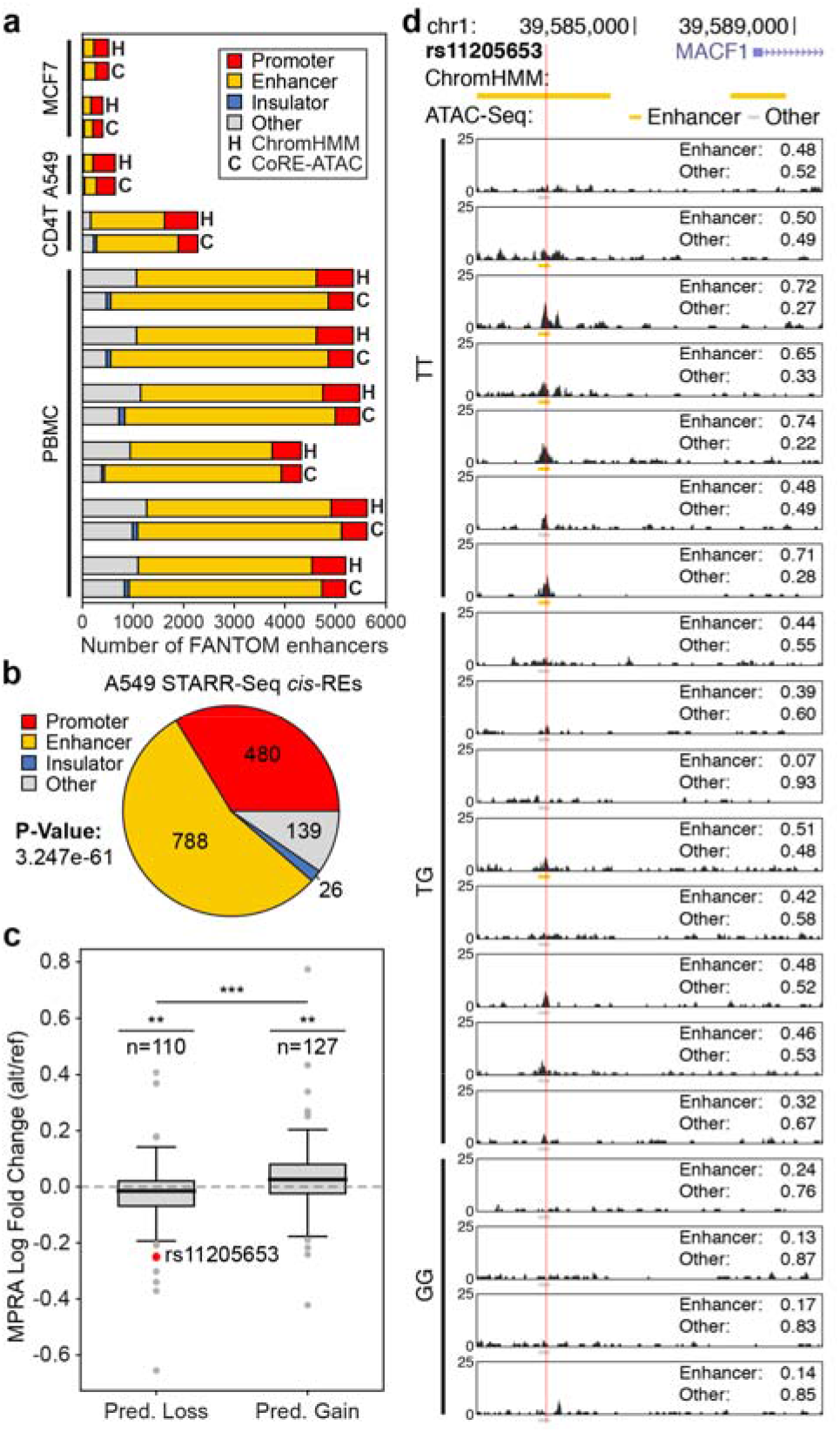
CoRE-ATAC predictions overlap with experimentally detected enhancers. (**a**) Overlap of FANTOM enhancer annotations with CoRE-ATAC (C) and ChromHMM (H) predictions in MCF7, A549, CD4^+^ T and PBMC samples. CoRE-ATAC predicted the majority of FANTOM enhancers as enhancers or promoters, recapitulating these experimentally identified enhancers. CoRE-ATAC annotations were similar to ChromHMM annotations. (**b**) CoRE-ATAC predictions for active regulatory regions identified by STARR-seq in A549 cell line. The majority of active enhancers identified by STARR-seq were predicted as promoter or enhancer by CoRE-ATAC. (**c**) MIN6 MPRA log fold change values for genomic regions predicted as losing or gaining RE function based on CoRE-ATAC probabilities for reference and alternative alleles. Significance for predicted loss and predicted gain categories was calculated using student’s t-test for MPRA log fold change values being less than or greater than 0 respectively. Significance comparing the predicted loss and predicted gain of MPRA fold change distributions was calculated using Mann-Whitney U test. We observed concordant direction of effect both for CoRE-ATAC predictions and MPRA acitivity levels when alternative and reference alleles are compared. (**d**) Genome browsers of 19 islet samples highlighting a loss of enhancer activity for rs11205653 (also highlighted in (**c**)) for the alternative allele (G). Values for enhancer and other represent the probability assigned to those classes of REs by CoRE-ATAC. We observe that for 5 out of 7 individuals with the reference allele (TT) CoRE-ATAC predicted enhancer activity, reflecting ChromHMM reference annotations, while for the individuals with GT or GG alleles, we observed an enhancer activity loss for all but one individual based on CoRE-ATAC predictions.

Another alternative method for investigating enhancer activity is massively parallel reporter assays (MPRA)[27], which enables experimentally testing thousands of sequences in parallel for regulatory activity (i.e., promoters and enhancers)[33]. To understand whether CoRE-ATAC could detect enhancers identified *via* MPRAs, we first compared enhancer predictions in A549 cells to enhancers identified from self-transcribing active regulatory region sequencing (STARR-seq)[26]. The majority of sequences that showed regulatory activity in this assay were predicted as enhancers (overlap significance calculated using Fisher’s exact test p-value = 3.247e-61) by CoRE-ATAC (Fig 4b). We noted that a significant portion of STARR-seq enhancers (~33%) were predicted as promoters in our models and the majority of these regions were close to TSS (S11 Fig a). In contrast, CoRE-ATAC enhancers overlapping STARR-seq active regions were more distal from the TSS (S11 Fig b).

Regulatory activities of certain open chromatin regions are genetically modulated, which can be detected *via* chromatin accessibility quantitative trait loci (caQTL) analyses. Previously, we identified caQTLs from human islet samples (n=19) [6] for which we generated an MPRA library to test and compare the regulatory activity of reference and alternative alleles for caQTLs and other variants (n=4293 SNPs) (manuscript in revision)[34]. These data gave us the opportunity to test whether CoRE-ATAC predictions can detect genetically driven differences in the regulatory activity. Using CoRE-ATAC prediction probabilities from 19 islets (stratified based on genotypes), we identified 237 loci for which a gain or loss of enhancer activity was predicted based on individuals’ genotypes and CoRE-ATAC prediction probabilities using one-tailed point-biserial correlation p-values (Methods). Among these, 110 loci lost activity in the alternative allele, whereas 127 gained activity. For these sequences, we compared the regulatory activity from MPRA assays to the predicted gain/loss of activity from CoRE-ATAC and confirmed that the direction of effect coincides with the two analyses (Fig 4c). More specifically, loci with gain of function for the alternative allele based on CoRE-ATAC predictions had higher MPRA activity for the alternative allele in comparison to the reference allele (i.e., fold change > 0). A similar concordance was observed for loci associated with loss of function for the alternative allele. For example, islet samples with the alternative allele for SNP rs11205653 (an islet caQTL) had lower enhancer probabilities compared to the samples with the reference allele, in agreement with the activity levels from the MPRA library for this locus (Fig 4d). Although CoRE-ATAC enhancer predictions were higher for individuals with TT genotype compared to individuals with GG and GT genotypes, we noted that individual-level heterogeneity in RE activity levels within the same genotype. This heterogeneity likely stems from non-genetic factors including disease status (diabetic versus healthy) or other clinical information (i.e., medication use, sex, race). Together, these results and findings establish CoRE-ATAC’s ability to predict individual level variability in *cis*-RE activity including heterogeneity stemming from genetic variation.

### Predicting disease-relevant enhancers from single nuclei ATAC-seq data

Single nucleus ATAC-seq (snATAC-seq) reveals chromatin accessibility at single nucleus resolution and enables 1) interrogation of chromatin accessibility at the single cell level; 2) identifying epigenomic maps of rare cell types *via* unsupervised clustering methods[14]; 3) enabling the study of cell types within tissues without the need to sort cells. Despite the benefit of single cell approaches and the shift in data generation from bulk to single cells, it is not established whether machine learning models built from bulk ATAC-seq data can be used for predictions in snATAC-seq data (either at the single cell or cell cluster level). To test this, we predicted *cis*-RE functions from human PBMC snATAC data[14] by first clustering cells into 15 groups based on the similarity of their accessibility profiles (Fig 5a). Comparisons with sorted immune cell bulk ATAC-seq data[15, 19] revealed 7 distinct cell types corresponding to B, natural killer (NK), CD8^+^ T, CD4^+^ T, effector CD4^+^ T, CD14^+^, and dendritic cells (DCs) (Fig 5b). CoRE-ATAC models trained on bulk ATAC-seq data, predicted *cis*-RE function in snATAC aggregate clusters (mean micro-average precision = 0.68) (Fig 5c), showing the flexibility and robustness of the method. Insulator predictions from snATAC-seq data were significantly enriched for CTCF/BORIS motifs (S12 Fig), confirming the biological relevance of these predictions.

**Fig 5.**
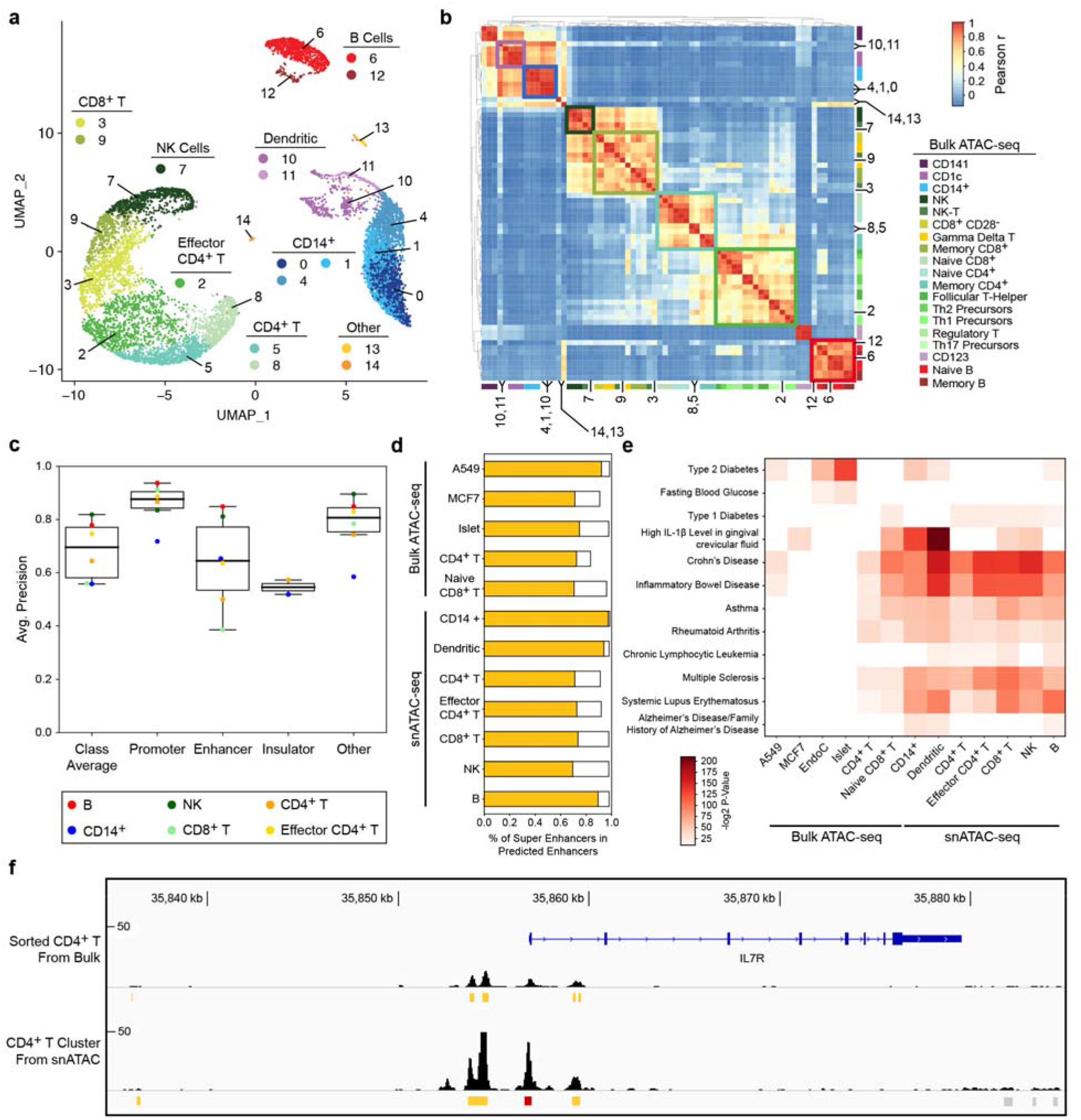
Predicting functionality of REs from clusters of PBMC snATAC-seq data. (**a**) Single cell clusters annotated for 7 immune cell types. Two-pass clustering identified a total of 15 cell clusters which we annotated using hierarchical clustering with sorted bulk ATAC-seq data (shown in (**b**)) to identify 7 different immune cells corresponding to these clusters. (**b**) Hierarchical clustering of scATAC clusters with bulk ATAC-seq data. Numbers and highlighted regions within the heatmap correspond to cell clusters and annotations in (**a**). 7 immune cell types were observed with both snATAC and bulk ATAC-seq samples. (**c**) Average precision values for predicting RE function in snATAC for the 7 annotated clusters. Model performances suggest that CoRE-ATAC is an effective tool for interrogating *cis*-RE activity from snATAC data. (**d**) Percent of super enhancers detected among CoRE-ATAC enhancers, demonstrating CoRE-ATAC’s ability to identify cell-type-specific enhancers that are most relevant to disease. (**e**) GREGOR SNP enrichment analysis highlighting selected diseases whose SNPs were significantly enriched within the enhancer elements predicted by CoRE-ATAC. Enhancers from snATAC-seq were significantly enriched for SNPs associated with immune diseases. (**f**) Genome browser view of *IL7R* for bulk ATAC and snATAC samples for CD4^+^ T cells. ATAC-seq read profiles and CoRE-ATAC predictions between snATAC and bulk ATAC were found to be similar to one another, demonstrating CoRE-ATAC as a robust method for *cis*-RE predictions. Red represents promoter predictions, yellow represent enhancer predictions, and gray represent “other” predictions from CnRE_-_ΔTΔC_

Similar to enhancer predictions in bulk ATAC-seq, the majority of CoRE-ATAC enhancer predictions in snATAC clusters were cell-specific (S13 Fig). To further establish CoRE-ATAC’s ability to identify disease relevant enhancers, we compared CoRE-ATAC predicted enhancers with super enhancers obtained from SEdb[35] in the 7 immune cell types. Super enhancers are cell-type-specific and harbor disease relevant SNPs[2]. On average, ~81% of super enhancers were captured by CoRE-ATAC enhancer predictions within their respective cell types for single cell predictions (Fig 5d), which was comparable to the detection rate from bulk ATAC-seq data which captured ~76% of super enhancers. The ability to identify cell-specific enhancers from snATAC-seq data is instrumental for studying rare cell populations and clinical samples.

We further assessed CoRE-ATAC’s ability to identify disease-relevant enhancers by conducting GWAS SNP enrichment analyses using GREGOR[36]. For this analysis we compared CoRE-ATAC enhancer prediction in the 7 cell types inferred from snATAC-seq data with enhancers predicted from bulk ATAC-seq data (A549, MCF7, islets, EndoC, CD4^+^ T and naïve CD8^+^ T). Immune cell enhancers predicted from snATAC-seq data were enriched in diseases related to the immune system (Fig 5e). For example, B cell enhancers were the most significantly enriched for variants linked to Systemic Lupus, a disease characterized by dysfunctions in B cells[37]. Islet enhancers were most significantly enriched for Type 2 Diabetes and Fasting Blood Glucose as expected, while T cell were most significantly enriched in immune diseases (e.g., Type 1 Diabetes). Overall, enhancer predictions were enriched in diseases that were the most relevant to their respective cell types, confirming that CoRE-ATAC can infer cell-specific enhancers.

Chromatin accessibility profiles of snATAC-seq clusters resembled that of bulk data from sorted cells, enabling CoRE-ATAC to effectively predict *cis*-RE function from snATAC-seq data, e.g., the locus around the *IL7R* (an important gene for T cell aging[19]) in CD4^+^ T cells both in bulk and single cell maps (Fig 5f). Despite similarities between sorted cell and single cell cluster epigenomes, the efficacy of CoRE-ATAC on snATAC-seq data was not given due to the differences in how cells are processed (i.e., FACS sorted cells versus single cell clusters) as well as differences between bulk and snATAC-seq libraries (depth, peak sizes, read distributions etc.). Our analyses have demonstrated that CoRE-ATAC models built from bulk data can effectively predict *cis*-RE function in snATAC-seq data, which is essential for the analyses of future snATAC-seq maps.

## Discussion

Recent advances in ATAC-seq profiling and single cell genomics revolutionized the epigenomics field by enabling the generation of chromatin accessibility maps from small starting material, even at single cell resolution. Due to these advances, epigenomic maps of human cells/tissues from many individuals are being generated at an unprecedented rate, including our own studies, to study how epigenomic landscapes change with age, diseases, and upon in-vivo and in-vitro activation [6, 12, 15–20]. These epigenomic maps are instrumental for inferring *cis*-REs from clinically relevant samples. CoRE-ATAC harnesses the power of deep learning to integrate chromatin accessibility maps with DNA sequence and effectively predicts the functionality of *cis*-REs (i.e., promoters, enhancers, insulators) across diverse cell types and individuals. We extensively evaluated CoRE-ATAC’s efficacy to predict *cis*-RE function across multiple cell types including those that are not used in model training. We established that CoRE-ATAC is an effective method for classifying the functional state of *cis*-REs, in new cell types (i.e., cell types that are missing reference annotations) and in snATAC-seq data.

CoRE-ATAC was trained using chromatin states identified using ChromHMM[11]. Despite ChromHMM being a computational method itself, it is a data driven approach that utilizes more data (i.e., multiple ChIP-seq assays) than a single ATAC-seq map to infer different chromatin states. The unsupervised approach of ChromHMM identifies clusters of genomic regions that have similar combinations of transcription factor/histone modification marks which can then be annotated using domain knowledge (e.g., states with H3K4me1, H3K27ac are likely enhancers). Although ChromHMM states are not directly and experimentally established *cis*-RE functions, they have multiple advantages over alternatives for class labeling in our machine learning models. First, ChromHMM profiles are genomewide providing many examples for model training. Second, ChromHMM states are available for many different cell types, enabling training and testing models across many cell/tissue types. Finally, ChromHMM states are well studied and functionally validated[38], making these annotations high-quality references, despite being computational inferences. Our method recapitulates ChromHMM annotations with a single assay (bulk or single nuclei ATAC-seq) as opposed to using multiple ChIP-seq assays, which enables the study of *cis*-REs in many cell types and tissues.

One of the unique features of CoRE-ATAC is its ability to integrate an individual’s genotype with chromatin accessibility maps by inferring the genotype from ATAC-seq read alignments. CoRE-ATAC accomplishes this with a sequence encoder that uses the frequency of ATAC-seq reads observing a specific base-pair at a genomic position instead of solely using a one hot encoding of the reference genome, separating our method from existing ones that use the commonly used one hot-encoding approach. If genotype data (e.g., SNP arrays) are available for an individual, one can also leverage this information in CoRE-ATAC predictions by providing a reference specific to the individual. Genotype-aware methods can be effective in predicting the annotation and activity of *cis*-REs whose activity is modulated by genetic variation (e.g., caQTLs[5–8]). In alignment with this, we demonstrated that CoRE-ATAC predictions from individual islet epigenomes aligned with enhancer loss/gain between alleles inferred from MPRA assays.

CoRE-ATAC also established a foundation for inferring functional annotations from snATAC-seq clusters, which can be useful to study rare cell populations that can be identified from these assays. For this analysis, we used CoRE-ATAC models trained on bulk data due to current limitations on the availability of snATAC-seq data that coincide with existing ChromHMM/*cis*-RE annotations. Although models from bulk data were effective in predicting functionality of *cis*-REs from single cell data, models built from snATAC-seq data might further improve the predictive performance. In the future, as more snATAC-seq data becomes available for more samples and more diverse cell types with established ChromHMM and CTCF insulator states, a CoRE-ATAC model can be trained solely on snATAC-seq data, potentially resulting in better classification performances for the purposes of annotating rare cell populations. Our predictions from snATAC-seq data were conducted at the cluster level. Given the sparsity of chromatin accessibility information at the individual single cell level (open in both alleles, open in one allele, closed), predicting functionality at the single cell level will remain a challenge.

CoRE-ATAC is widely applicable for inferring *cis*-RE function from ATAC-seq and further improves upon naïve methods as well as existing machine-learning models (Fig 2d, Fig 3b) through its ability to leverage ATAC-seq signal information. Although model training is a computationally expensive process, requiring GPUs to train in a timely manner, using these models for predictions is much faster, requiring a little over 2 minutes (excluding the time to load the data into memory) to functionally annotate *cis*-REs for 75000 loci using a 2.3 GHz 8-Core Intel Core i9 processor (Fig S14). Our indepth performance analyses suggest that CoRE-ATAC can be widely used, even with limited computational resources, to improve the functional annotations of ATAC-seq *cis*-RE maps. To promote the widespread use of our predictive model we have made the CoRE-ATAC code and pre-trained model freely available on GitHub (https://github.com/UcarLab/CoRE-ATAC).

## Materials and Methods

### Machine Learning Architecture

CoRE-ATAC utilizes both data encoded in the deep learning framework and features extracted using our previous method PEAS[21]. These two feature sets are provided as two separate inputs into a machine learning model implemented using Keras[39] and Tensorflow[40] libraries. The deep learning component uses four convolutional layer blocks. Each block consists of i) a 1D convolutional neural network (CNN) layer (window size = 19), ii) a batch normalization layer, iii) a second 1D CNN layer (window size = 19) iv) a max pooling layer (pool size =2) and v) a second batch normalization layer, in this order (Fig 1). The first two blocks of convolutional layer units utilize 256 filters while the final two blocks utilize 512 filters. Both the deep learning component and the PEAS component of the model are trained using their own dense neural network layer with n=2048 nodes. The output of these dense layers is then concatenated and provided as input into a combined dense layer with n=512 nodes before classifying *cis*-REs in the final output dense layer using the Adam[41] optimization method. CNN layer window sizes, number of filters, and number of dense layer nodes were selected based on the best parameters observed during model tuning.

### ATAC-seq Data Encoders

We implemented a novel ATAC-seq data encoder that takes as input: paired-end reads, peaks, and the reference genome (or a personalized genome reflecting an individual’s genetic variation) to generate one 10×600 matrix per peak. Rows represent DNA sequence and ATAC-seq signal data, while columns represent the 300 base-pair positions downstream and upstream from the peak center (Fig 1). Details of this matrix are explained below:

#### Rows 1-4

The 0-1 normalized frequency of observing 1) adenine (A), 2) cytosine (C), 3) guanine (G), or 4) thymine (T) from ATAC-seq read pileups, which is calculated when there is enough read coverage (n >= 10). For low coverage loci; the frequency reduces to a one-hot encoding, representing the corresponding nucleotide in the reference genome.

#### Row 5

The number of insert pileups within the 600 base-pair window, where each position is z-standardized with respect to pileups across all peaks for that position.

#### Rows 6 and 7

The number of 5’ (row 6) and 3’ (row 7) cuts at a given position (z-standardized). Savitzky-Golay smoothing filter[42] is applied with a window size of 15 to account for an increase in the number of cuts across multiple positions in close proximity due to sequencing depth.

#### Rows 8 and 9

The median fragment length for 5’ and 3’ cut site, applying the same standardization and smoothing used for rows 6/7.

#### Row 10

The original peak region, setting values to 1 if a base-pair position is within the peak and 0 otherwise. This allows for distinguishing between peaks that are in close proximity to one another and providing an indicator of the most relevant positions.

In all analyses, peaks used for data encoding were called with MACS2[43] using parameters “-f BAMPE --nomodel”. Duplicates were kept only in the snATAC-seq data analyses using the “--keep-dup all” option.

### Ground Truth Selection & Model Training

CoRE-ATAC was trained on ATAC-seq data from 4 different cell types: GM12878[12] (GSE47753), K562[20] (GSE121993), HSMM[18] (GSE109828, donor “54-1” at time 0), and CD14^+^ monocytes[19] (EGAS00001002605) (Table 1). Class labels were assigned by co-analyzing 15-state ChromHMM[11] models generated in these cell types with corresponding 18-state Roadmap[10] ChromHMM annotations. 15 state ChromHMM models were trained for each cell type independently using H3K4me1, H3K4me3, H3K27ac, H3K27me3, H3K36me3, H3K9me3 and CTCF ChIP-seq data from ENCODE[9]. Hierarchical clustering of emission probabilities revealed 10 distinct clusters (S1 Fig) corresponding to active promoters, promoters, flanking enhancers, active enhancers, enhancers, genic enhancers, transcribed, insulator, repressed, and low signal states. Preliminary machine learning models were trained using the 10 clusters identified and peaks were selected based on their concordance with Roadmap 18 state models (S2 Fig) in order to select the best examples for model training. Performance evaluations on the 10-state models revealed that only 4 major classes of *cis*-REs could be effectively predicted from such models across all cell types: promoter, enhancer, insulator and other classes (S3 Fig). Based on these results, we focused on training models with these 4 major classes and selected peaks using a wider range of concordance with Roadmap states (S4 Fig).

To make CoRE-ATAC predictions robust to instances of low chromatin accessibility levels or false positives in ATAC-seq peak calls, non-peak locations were also included for a subset of samples. We selected non-peak locations by taking the union of all peaks across all 6 samples and selecting additional genomic regions for HSMM, CD14^+^ sample 2 (S2), and K562 replicate 2 (R2) samples that were called as peaks in other samples but were not called within the sample itself. Only half of the samples were selected to ensure that CoRE-ATAC could be robust to differences in normalization steps when provided either a list of genomic regions that could contain no ATAC-seq reads or when provided ATAC-seq peaks where every region must contain read information to call the peak at the given location. We further filtered these regions to ensure that ChromHMM of these additional annotations were annotated as “other” in the respective cell types. Finally, we randomly selected non-peak regions so that only up to 25% of the “other” category for the sample was represented by these non-peak regions. This additional filtering was applied to ensure that non-peak regions did not dominate the “other” class to the point of CoRE-ATAC only classifying peaks as “other” if the genomic regions have reduced or no ATAC-seq read counts. In total, 153311 loci across 4 cell types were used in model training for 4-state classification, with 2551 of these loci representing non-peak regions.

Deep learning and PEAS components of CoRE-ATAC were initially trained separately to handle different rates of overfitting observed between the two models (overfitting evaluated using training and validation loss). Parameter tuning was performed in this stage of training to adjust i) the number of filters, ii) convolutional layer window size, iii) number of convolutional blocks, and iv) number of nodes within dense layers (where applicable) using held out validation data (i.e., ATAC-seq peaks in chromosomes 2 and 10) (Table 2). Individually each component achieved 82.56% and 79.25% accuracy on held out validation data for deep learning and PEAS components respectively using the best observed parameters for minimizing the validation loss. We then combined these models into a single unified model by concatenating deep learning and PEAS components with a concatenation layer, for a final round of model training.

### Model Tuning

Hyperparameter selection was performed on i) the 1D convolution kernel size {9,11,19, 21}, ii) the number of convolutional layer block {3,4,5}, iii) intermediate dense layer size {1024, 2048, 4096} and iv) and the number of convolutional layer filters {32,128, 256} where for each block the first two convolutional layers were set to the initial filter size and then doubled for the remaining two convolutional layers. Due to the long training time on these models, parameter, not every combination (108 total) was tested, instead different parameters were tested one by one, and the best performing value for the parameter was kept.

### Model Evaluation

To test the performance of CoRE-ATAC on datasets and cell types that were not used in model training, we predicted *cis*-REs in ATAC-seq data obtained for 7 different cell types: MCF7[16] (GSE97583) (n=2), Naïve CD8^+^ T[15,19] (GSE118189 & EGAS00001002605) (n=10), Peripheral Blood Mononuclear Cells (PBMCs) [15, 19] (EGAS00001002605) (n=6), CD4^+^ T[12] (GSE47753) (n=1), A549[44] (GSE117089) (n=1), pancreatic islets[6] (SRP117935) (n=19) and EndoC beta cell line[17] (GSE118588) (n=1) (Table 1). Model performances were evaluated using ChromHMM states in each cell type.

### Annotations for Model Evaluation

ChromHMM 18 state model annotations from Roadmap[10] were used for naïve CD8^+^ T and A549, where annotations were converted to 4 major classes (promoter, enhancer, insulator and other), mapping states to the most relevant class label. “TSSA” and “TssFlnk” were mapped to promoter, “EnhG1”, “EnhG2”, “EnhA1”, “EnhA2”, and “EnhWk” were mapped to enhancer and “Tx”, “TxWk”, “ZNF/Rpts”, “Het”, “ReprPC”, “ReprPCWk”, and “Quies” were mapped to “other”. Peaks that could not be mapped to these major classes (i.e., “TssFlnkU”, “TSSFlnkD”, “TSSBiv”, and “EnhBiv”) were excluded. For PBMC and CD4^+^ T samples, 15 state ChromHMM models were trained using H3K4me1, H3K4me3, H3K27ac, H3K27me3, H3K36me3, and H3K9me3, ChIP-seq from Roadmap. ChromHMM states for MCF7, Islet, and EndoC were obtained from the following studies[3, 17] and were converted to the 4 major class labels used by CoRE-ATAC. Only ATAC-seq peaks with greater than 90% overlap with a single class label were kept to filter out ambiguous regions.

### Comparison with alternative methods/assays

We compared CoRE-ATAC to two sequence-based methods (DeepSEA[24] and LS-GKM[28]) and our previous neural network (NN) based method (PEAS[21]). For this comparison we focused on enhancer versus “other” predictions, the most difficult discrimination task in our models (Fig 2c). For this, we chose all enhancers and “other” annotated regions from our ground truth test set (i.e., regions within chromosomes 3 and 11 for GM12878, K562, HSMM, and CD14^+^ samples) (Table 1 and Table 2). The same training and testing set was used for CoRE-ATAC, PEAS, and LS-GKM. For DeepSEA, we used the web annotation tool (http://deepsea.princeton.edu). DeepSEA makes multiple predictions of activity for a wide array histone marks and transcription factors (TFs) across multiple cell types. We therefore selected enhancer probabilities by taking the maximum probability score for H3K4me1 and H3K27ac across all cell types predicted by DeepSEA. The area under the receiver operating characteristic curve (ROC AUC) and average precision metrics were used to compare the four methods (Fig 2d, Fig S6).

Naïve method comparisons focused on the 40 samples not used in model training to fairly assess CoRE-ATAC’s performance with respect to these methods. Multiple thresholds were applied for each Naïve method to identify the best threshold to set for predicting promoters, enhancers, and insulators. Promoters were tested using 1kb, 2kb, and 5kb distances to the nearest TSS, (distances calculated using HOMER[29]). Enhancers were tested using MACS2[43] FDR qval of 0.01, 0.001, and 0.0001.

Insulators were tested using the number of CTCF motifs greater than 0, 2, and 4. Promoters, enhancers, and insulators were selected using these thresholds, assigning probabilities 1.0 when the threshold requirement is met, and 0.0 otherwise. Finally, combined naïve approach performance was calculated selecting the best threshold (i.e., 2kb, 0.001, and 0 for promoter TSS, enhancer MACS2 qval, and insulator number of CTCF motifs respectively). Due to the nature of selecting enhancers using MACS2 qval, priority was given to promoters and then insulators. For the remaining regions that were not annotated as either promoter or insulator, enhancers and “other” were classified using MACS2 qval. Performances were evaluated using Matthews correlation coefficient.

Alternative enhancer definitions were explored to understand how well CoRE-ATAC can predict active regulatory elements (i.e., promoter and enhancers) from FANTOM[31, 32] and STARR-seq[26]. FANTOM enhancers identified using Cap Analysis of Gene Expression (CAGE) technology[25] were obtained for MCF7, A549, CD4^+^ T cells, and PBMCs. We obtained STARR-seq active regulatory sites for A549[45] from GEO[46, 47] (GSE114063). These regions were then compared with CoRE-ATAC predictions, counting the number of predictions for each class within FANTOM5 enhancers and STARR-seq sites. Fisher’s exact test was used to calculate the significance of CoRE-ATAC enhancers overlapping with Starr-seq enhancers.

Massively Parallel Reporter Assay (MPRA) data were generated in MIN6 pancreatic beta cell line to study the regulatory activity of variants associated with Type-2-Diabetes (manuscript in revision) [34]. Briefly, 4293 variants within islet ATAC-seq peaks were profiled for regulatory element activity using MPRA. Taking the union of islet peaks called across all 19 samples, we predicted *cis*-RE functions for each islet sample to obtain class probabilities for each region and islet sample. We then putatively identified *cis*-REs showing loss or gain of *cis*-RE activity using one-tailed point-biserial correlation p-values, identifying loci with probabilities that were significantly lower or higher between reference and alternative genotypes. The maximum absolute value correlation was obtained for each peak with genotype information by calculating the correlations for all comparisons among ref/ref, ref/alt, and alt/alt genotypes, using the sum of promoter and enhancer probabilities. Peaks with significant point-biserial correlation coefficients (p-value < 0.01) were separated into two groups corresponding to the loss or gain of *cis*-RE activity (negative and positive correlations respectively) based on CoRE-ATAC predictions for different alleles. Finally, MPRA activity differences between alternative and reference alleles (log fold change in MPRA analyses) were compared with the activity differences inferred from CoRE-ATAC *cis*-RE class probabilities. Student t-test, and Mann Whitney U test were used to calculate the significance of MPRA log fold change values observed for predicted loss and gain of *cis*-RE activity both individually as well as comparatively.

### snATAC seq data analyses and clustering

Single nuclei ATAC-seq (snATAC) PBMC data[14] was obtained from GEO[46, 47] (GSE129785). Sequence reads were processed using Cell Ranger and cells were clustered using our own implementation of a recently described snATAC clustering method[14], which uses two passes of clustering to identify cell type clusters. In the first pass, genomic regions were binned into 2500bp windows, counting the number of paired-end reads within each bin. The top 50,000 bins with the greatest number of reads were selected for clustering using Seurat[48, 49], requiring a minimum of 185 cells per cluster. Peaks were called for each cluster independently using MACS2 (“-f BAMPE -nomodel, --keep-dup all”) and were used to perform a second pass of cell clustering as before, using peaks instead of bins, identifying a total of 15 clusters.

Cell type annotations for these clusters were obtained by comparing snATAC profiles with ATAC-seq profiles of sorted immune cell types *via* flow cytometry. To identify marker peaks for each cell type, we selected 2-3 representative and high quality ATAC-seq samples per cell type[15, 19] for 19 different immune cell types based on ATAC-seq library quality (FRIP score and read depth) and similarity between biological replicates (Spearman r). Marker peaks were identified from the signature profile generated by CIBERSORT[50], using its data pre-processing step. Hierarchical clustering was performed for read pileups within marker peaks for snATAC-seq clusters and sorted bulk data, which enabled annotating 15 clusters into 7 cell types. Cells that belong to the same cell type were pooled together and the functionality of *cis*-REs were predicted using CoRE-ATAC for 7 cell types by allowing duplicate reads. Predictions were evaluated using either ChromHMM annotations from Roadmap, or in-house ChromHMM states (CD4^+^ T and CD14^+^ monocytes).

### SNP enrichments

GWAS SNP enrichments were performed with GREGOR[36] software using index SNPs as well as linked SNPs (linkage disequilibrium threshold of R^2^ 0.7 for the European (EUR) population). NHGRI-EBI GWAS Catalog SNPs[51] (Obtained January 8^th^ 2020) for 3981 traits/diseases were used in enrichment analyses for different enhancer sets inferred by CoRE-ATAC.

### Super enhancer analysis

Super enhancer annotations were obtained for A549, MCF7, Islets, CD14^+^, CD4^+^ T, CD8^+^ T, monocyte-derived dendritic cells, NK, and B cells from SEdb[35]. We then calculated the percent of super enhancers found among CoRE-ATAC enhancer predictions to measure how well CoRE-ATAC predictions recapitulate these cell-type-specific enhancers.

## Declarations

### Ethics approval and consent to participate

Not applicable

### Consent for publication

Not applicable

### Availability of data and materials

The publicly available datasets analyzed during the current study are available from the GEO and EGA repositories and websites: GM12878 & CD4T ATAC-seq: https://www.ncbi.nlm.nih.gov/geo/query/acc.cgi?acc=GSE47753

K562 ATAC-seq: https://www.ncbi.nlm.nih.gov/geo/query/acc.cgi?acc=GSE121993

HSMM ATAC-seq: https://www.ncbi.nlm.nih.gov/geo/query/acc.cgi?acc=GSE109828

MCF7 ATAC-seq: https://www.ncbi.nlm.nih.gov/geo/query/acc.cgi?acc=GSE97583

Naive CD8 ATAC-seq: https://www.ncbi.nlm.nih.gov/geo/query/acc.cgi?acc=GSE118189 https://ega-archive.org/studies/EGAS00001002605

PBMC ATAC-seq: https://ega-archive.org/studies/EGAS00001002605

A549 ATAC-seq: https://www.ncbi.nlm.nih.gov/geo/query/acc.cgi?acc=GSE117089

EndoC ATAC-seq: https://www.ncbi.nlm.nih.gov/geo/query/acc.cgi?acc=GSE118588

A549 Starr-seq: https://www.ncbi.nlm.nih.gov/geo/query/acc.cgi?acc=GSE114063

snATAC PBMC data: https://www.ncbi.nlm.nih.gov/geo/query/acc.cgi?acc=GSE129785

FANTOM enhancers: https://fantom.gsc.riken.jp/5/datafiles/latest/extra/Enhancers/

Encode ChIP-seq: https://www.encodeproject.org/

Roadmap ChromHMM states: https://egg2.wustl.edu/roadmap/webportal/chr_state_learning.html

Islet MIN6 MPRA data analyzed during the current study are not publicly available yet due as the publication for these data is currently under revision, but are available from the corresponding author on reasonable request.

CoRE-ATAC code and pretrained models are available on our GitHub page: https://github.com/UcarLab/CoRE-ATAC

### Competing Interests

The authors declare that they have no competing interests

### Funding

This research was supported by the PhRMA Foundation postdoctoral fellowship award in bioinformatics (to AT) and National Institute of General Medical Sciences (NIGMS) under award number GM124922 (to DU).

### Author’s Contributions

AT and DU designed the study. AT implemented the CoRE-ATAC method, trained the deep learning model and conducted all downstream analyses. SK, RT, and MLS provided the MIN6 MPRA data and additional support for analyses conducted using these data. AT and AE processed the snATAC-seq data performed snATAC clustering of PBMCs. AT and DU annotated the final snATAC clusters. AT and DU wrote the manuscript. All authors read and approved the final manuscript.

## Acknowledgements

We thank members of the Ucar, Stitzel, Beck, and Lee labs at JAX-GM for their insightful feedback throughout this project.

## Notes

https://github.com/UcarLab/CoRE-ATAC/

### Competing Interest Statement

The authors have declared no competing interest.

