## Supplementary Figures for "CoRE-ATAC: A deep learning model for the functional classification of regulatory elements from single cell and bulk ATAC-seq data"

### Supplementary Figure 1

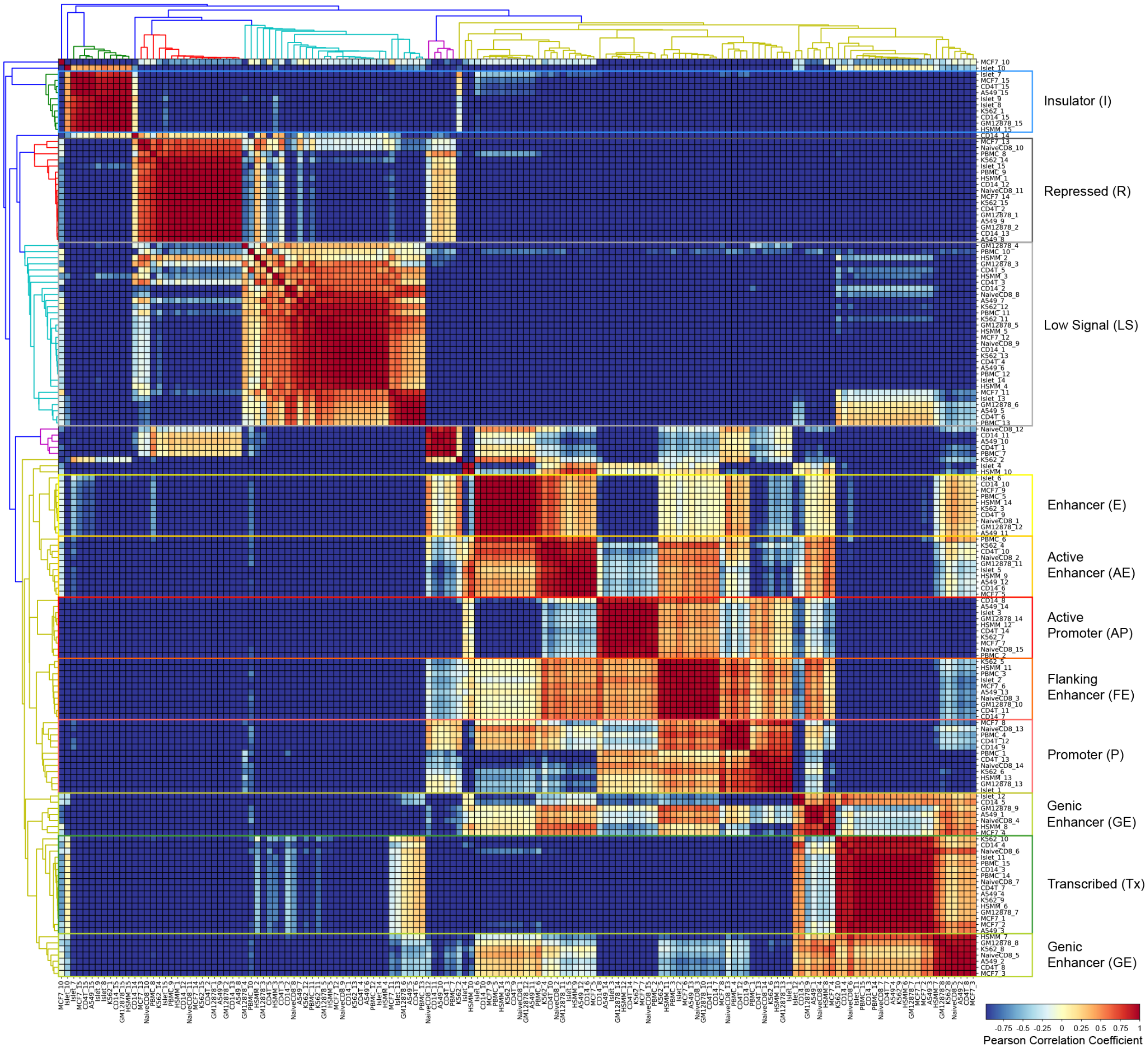


**S1 Fig. ChromHMM emission probability clusters reveal 10 consistent chromatin states.** We clustered the ChromHMM emission probabilities using Pearson's correlation coefficient to identify 10 chromatin states that were consistently present among all 10 cell types for which we recalled ChromHMM states. These 10 functional states include, insulators, repressed, low signal, enhancer, active enhancers, active promoter, flanking enhancer, promoter, genic enhancer, and transcribed regions. Numbers preceding the cell types are the emission states corresponding to Fig. S4. Note that genic enhancers are shown by two groups which correspond to active genic enhancers and genic enhancers which we combined due to the low number of cell types included within these clusters independently. Also note that for insulators, we selected islet state 7 as this was the state with the strongest CTCF signal (S4 Fig), maintaining one insulator state per cell type.

### Supplementary Figure 2

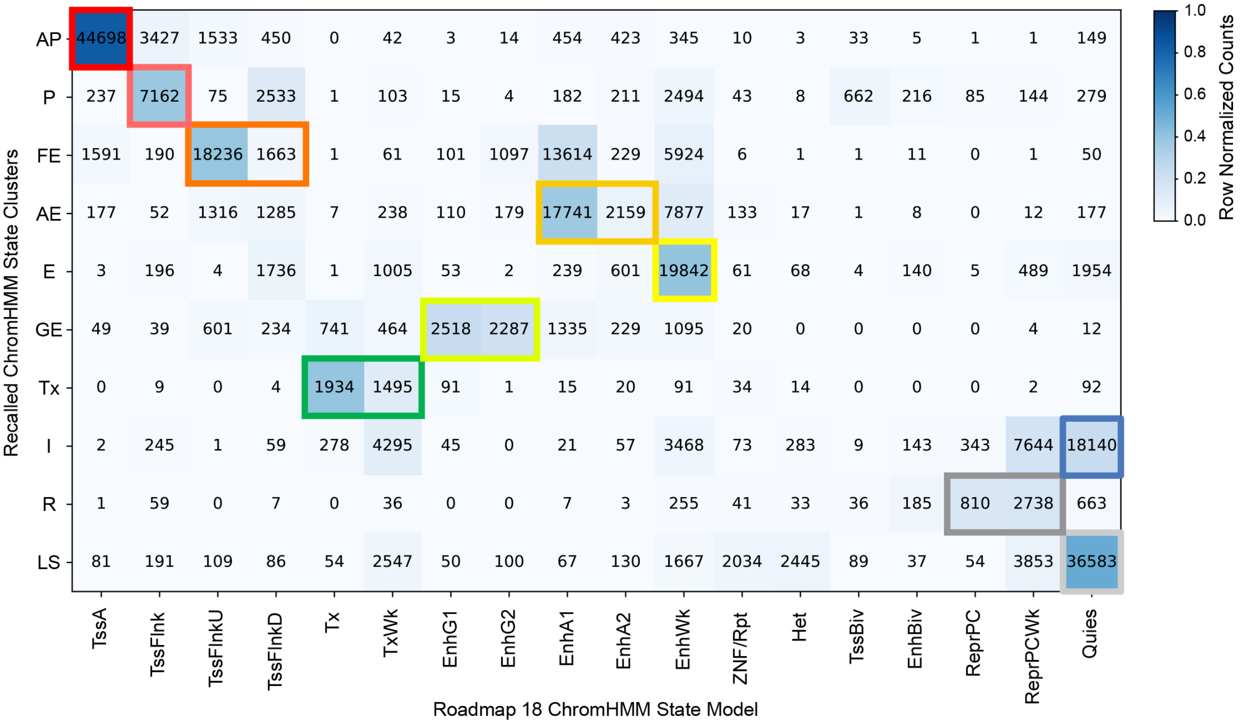


| | GM12878 | HSMM | K562 R1 | K562 R2 | CD14⁺ S1 | CD14⁺ S2 |
|---|---|---|---|---|---|---|
| **Active Promoter (AP)** | 11572 | 6753 | 7550 | 7234 | 5876 | 5713 |
| **Promoter (P)** | 2086 | 1766 | 1239 | 1211 | 433 | 427 |
| **Flanking Enhancer (FE)** | 8592 | 1192 | 3767 | 3539 | 1477 | 1332 |
| **Active Enhancer (AE)** | 3505 | 9720 | 1246 | 1212 | 2133 | 2084 |
| **Enhancer (E)** | 7989 | 4998 | 1711 | 1825 | 1853 | 1466 |
| **Genic Enhancer (GE)** | 2078 | 1192 | 500 | 485 | 282 | 268 |
| **Transcribed (Tx)** | 1478 | 493 | 303 | 370 | 358 | 427 |
| **Insulator (I)** | 9963 | 1367 | 1558 | 1324 | 2217 | 1711 |
| **Repressed (R)** | 1629 | 393 | 377 | 507 | 330 | 312 |
| **Low Signal (LS)** | 26845 | 4050 | 1940 | 2214 | 717 | 817 |

**S2 Fig. CoRE-ATAC 10 State model ground truth selection.** To select a ground truth for predicting the 10 functional states we identified, we corroborated our ChromHMM state calls with Roadmap 18 state models. (Top) Highlighted concordant recalled ChromHMM states with corresponding Roadmaps states to select active promoters (red), promoters (pink), flanking enhancers (orange), active enhancers (orange yellow), enhancers (yellow), genic enhancers (greenish yellow), transcribed (dark green), insulator (blue), repressed (dark gray), and low signal (light gray) functional states. (Bottom) The number of ground truth examples for each cell type used in model training and functional state examples selected.

### Supplementary Figure 3

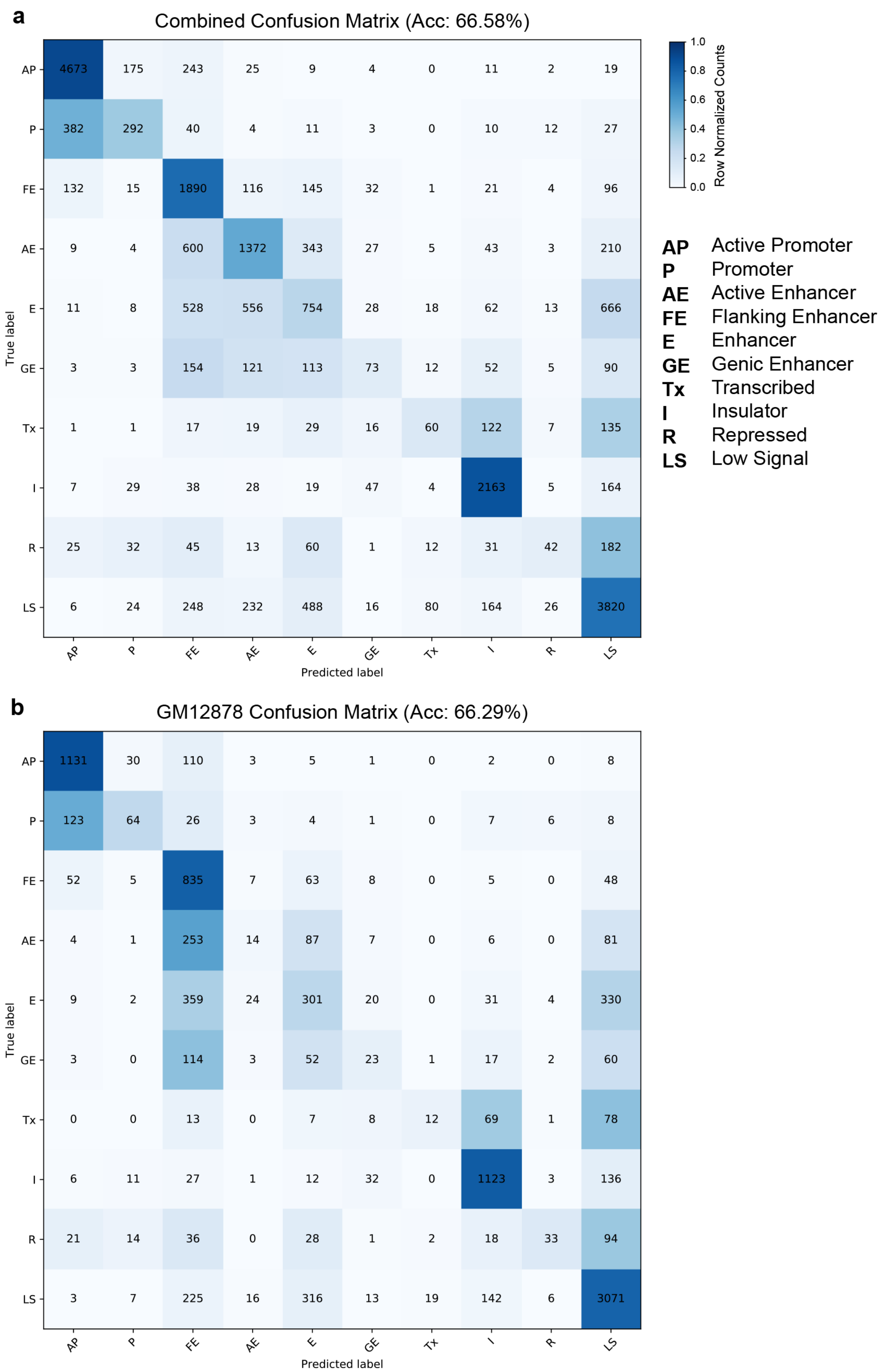


**S3 Fig. CoRE-ATAC 10 State performances.** (**a**) Confusion matrices of combined CoRE-ATAC performances for 10 state models. (**b**) Confusion matrices of CoRE-ATAC performance for 10 state models in GM12878. CoRE-ATAC 10-state models. Only 4 of the 10 chromatin states were predicted by CoRE-ATAC. Smaller subsets of these four functional states (i.e., promoters, enhancers, insulators and other) were predicted as the state with the highest number of examples.

### Supplementary Figure 4

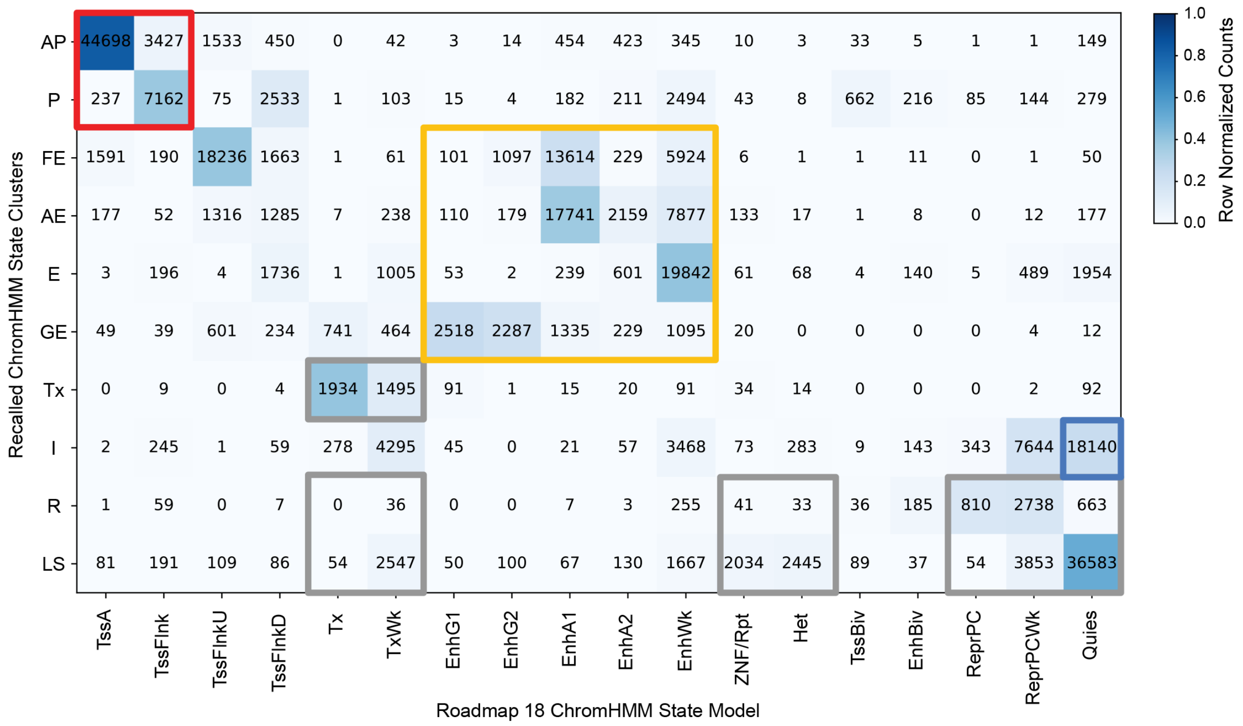


| | GM12878 | HSMM | K562 R1 | K562 R2 | CD14+ S1 | CD14+ S2 |
|---|---|---|---|---|---|---|
| **Promoter (P)** | 13883 | 8997 | 9020 | 8685 | 7550 | 7389 |
| **Enhancer (E)** | 16103 | 19133 | 5434 | 5442 | 16395 | 14725 |
| **Insulator (I)** | 9963 | 1367 | 1558 | 1324 | 2217 | 1711 |
| **Other (O)** | 36928 | 6500 | 3257 | 3823 | 2293 | 2519 |
| **Additional Non-Peaks** | 0 | 388 | 0 | 1315 | 0 | 848 |

**S4 Fig. CoRE-ATAC 4 State model ground truth selection.** (Top) Highlighted concordant recalled ChromHMM states with corresponding Roadmaps states to select promoters, enhancers, insulators and other. (Bottom) The number of ground truth examples for each cell type used in model training and functional state examples selected. Flanking enhancers were excluded as these regions are ambiguous and could be annotated as promoters. Merging states and relaxing concordance with Roadmap allowed for selecting more examples for model training.

### Supplementary Figure 5

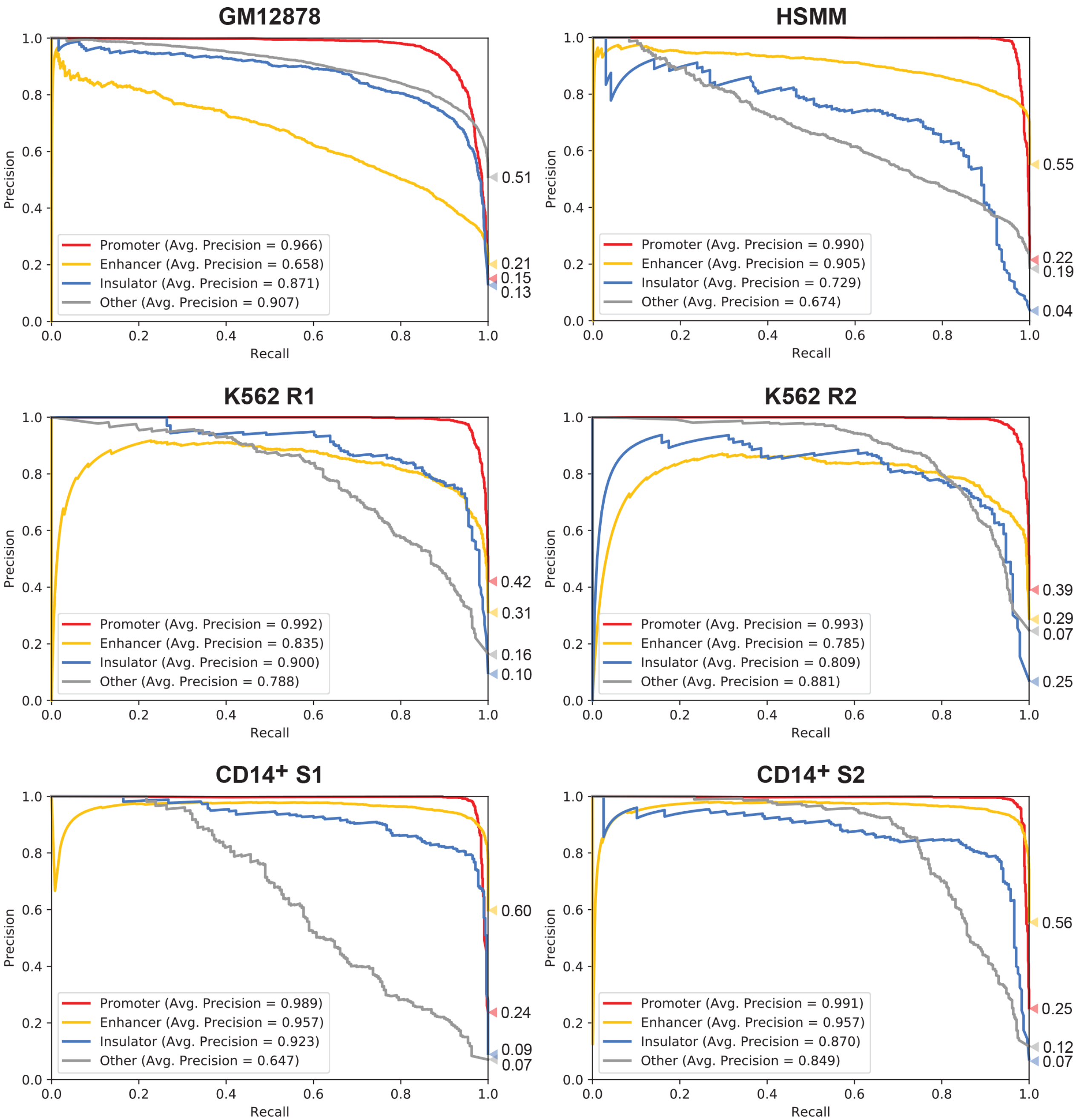


**S5 Fig. CoRE-ATAC predicts all functional annotations with high precision.** Precision recall curves of held out test data for each sample used in model training. Individual class performances reveal that CoRE-ATAC predicts all classes with high average precision.

### Supplementary Figure 6

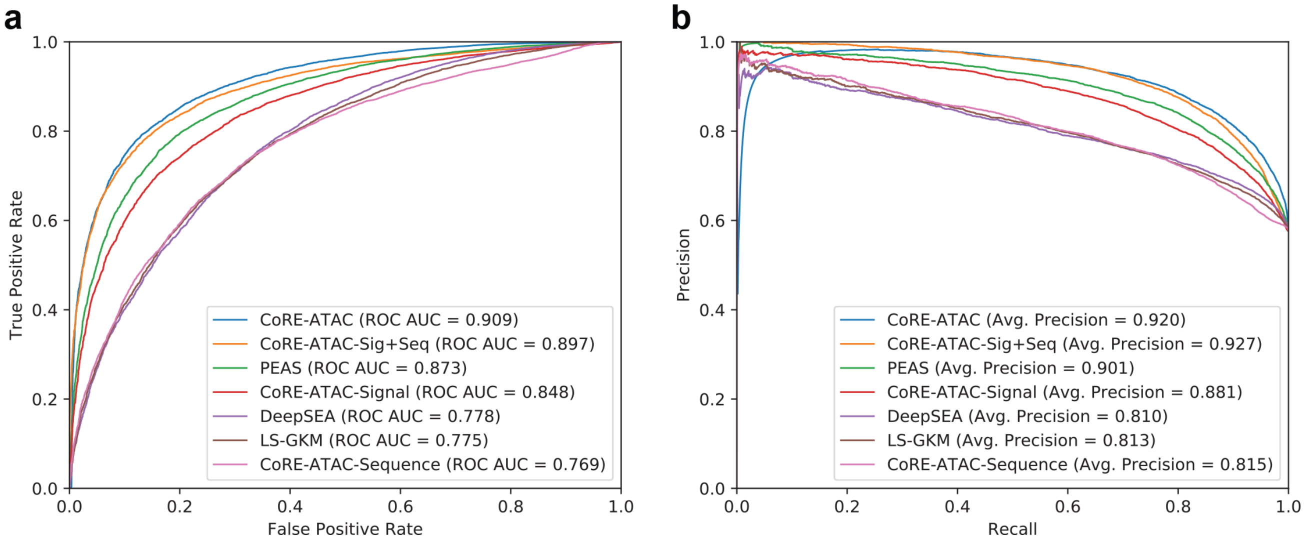


**S6 Fig. CoRE-ATAC component comparisons with sequence-based enhancer predictions.** Receiver operating characteristic (ROC) curves (**a**) and Precision Recall curves (**b**) for different enhancer prediction models: CoRE-ATAC components, PEAS, DeepSEA and LS-GKM. Sequence based approaches had similar performances, including CoRE-ATAC's sequence-based component. The ATAC-seq signal based (CoRE-ATAC-Signal) component of CoRE-ATAC alone outperforms all sequence-based approaches, however combining both sequence and signal greatly enhances predictive performances. PEAS captures more information than signal alone, however, is still underperforming compared to CoRE-ATAC-Sig+Seq model. Finally, the CoRE-ATAC model, which includes PEAS features, showed a slight improvement over the CoRE-ATAC-Sig+Seq model, likely taking advantage of the known features such as number of known motifs and conservation scores used in PEAS.

### Supplementary Figure 7

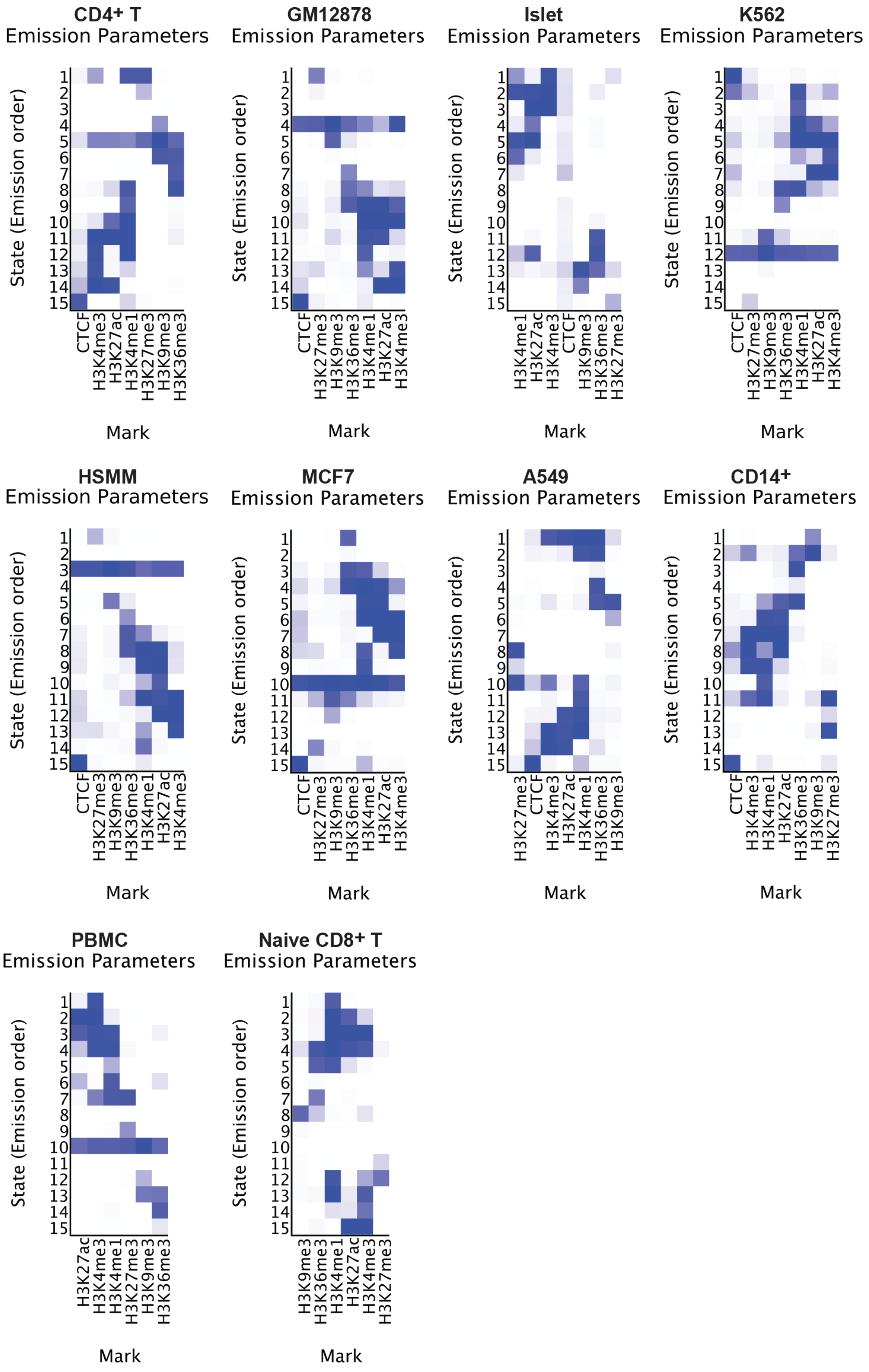


**S7 Fig. ChromHMM emission probabilities for all cell types.** Recalled ChromHMM states for 10 of the 11 cell types used in this study. Each chromHMM run included enhancer marker H3k4me1, promoter marker H3k4me3, repressor marker H3k27me3, active *cis*-RE marker H3k27ac, transcribed marker H3k36me3, heterochromatin marker H3k9me3, and CTCF insulator marker CTCF (when available). Emission probabilities revealed consistent histone modification mark combinations present throughout these diverse cell types.

### Supplementary Figure 8

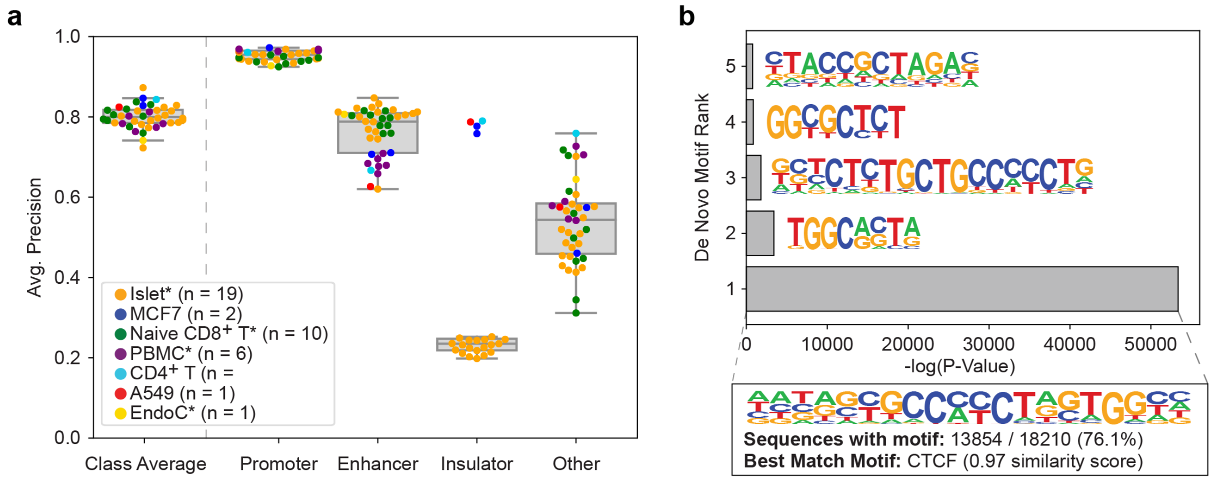


**S8 Fig. CoRE-ATAC Islet insulator predictions.** (**a**) Cross cell type model performances when including islet insulators. Using poor quality CTCF ChIP-seq data (as evident from S4 Fig) as a ground truth resulted in reduced model performances for all islet samples. (**b**) De novo motif enrichment for CoRE-ATAC insulator predictions in islets. Predicted insulators are highly enriched for CTCF with 76.1% of regions harboring a CTCF motif. Incorporating both DNA sequence and ATAC-seq signal enables CoRE-ATAC to detect a majority of insulators using the CTCF motif while detecting other insulators *via* other features learned in model training.

**Supplementary Figure 9**

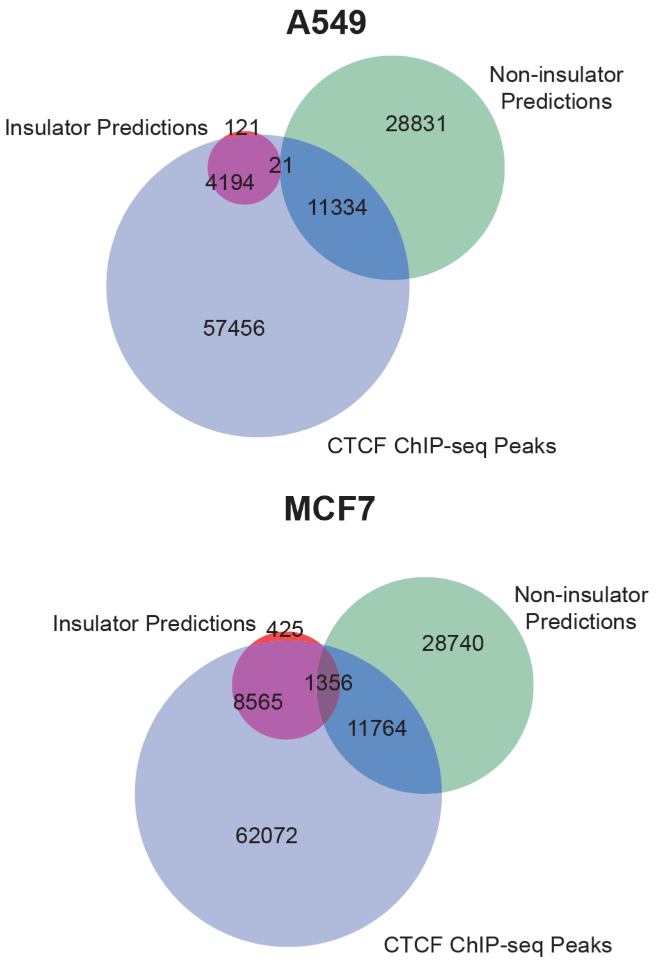


**S9 Fig. CTCF enrichment of CoRE-ATAC insulator predictions.** Overlap of CTCF ChIP-seq peaks with CoRE-ATAC insulator and non-insulator predictions. Majority of CoRE-ATAC insulator predictions overlapped with CTCF ChIP-seq peaks.

### Supplementary Figure 10

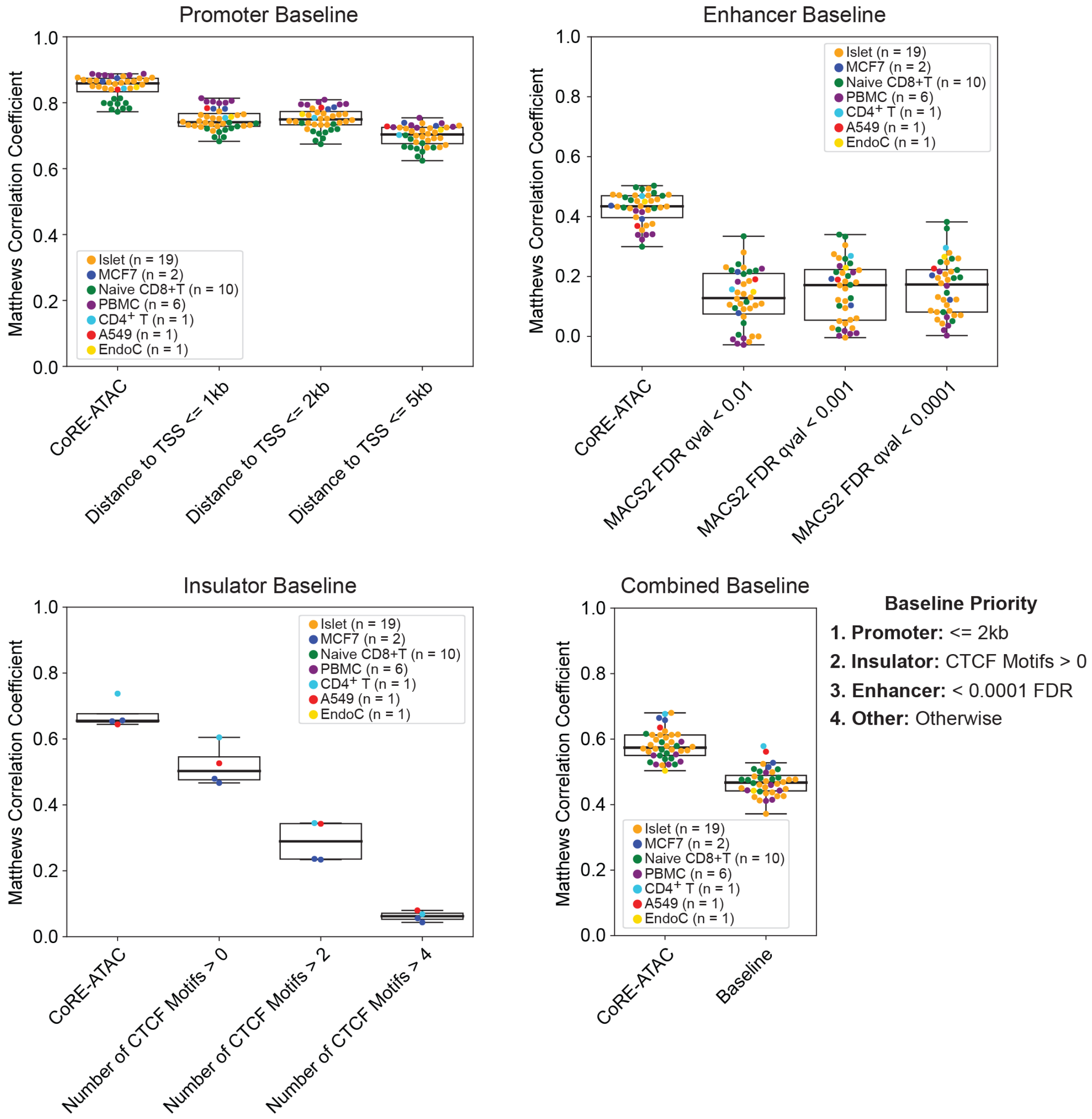


**S10 Fig. CoRE-ATAC outperforms threshold-based naïve/baseline annotations.** CoRE-ATAC outperforms each threshold-based method for detecting promoters, enhancers, insulators, and combined annotations. Matthews Correlation Coefficient was used to measure performances as it accounts for true positives, true negatives, false positives, and false negatives simultaneously within its function and is an ideal measurement for performances when probabilities are static as in the case with all threshold-based approaches (i.e., probability = 1.0 if satisfying the threshold, 0.0 otherwise). CoRE-ATAC consistently had the best performance compared to each baseline. For detecting promoters, the best threshold was identified as ATAC-seq peaks within 2kb of the promoter, for enhancers, an FDR qval score is less than 0.0001, and for insulators, whether or not the region contained a single (n=1) CTCF motif was the best performing threshold. Finally, combining all three of these thresholds confirmed that CoRE-ATAC improves our ability to predict *cis*-RE function, outperforming commonly used methods for detecting promoters, enhancers and insulators.

### Supplementary Figure 11

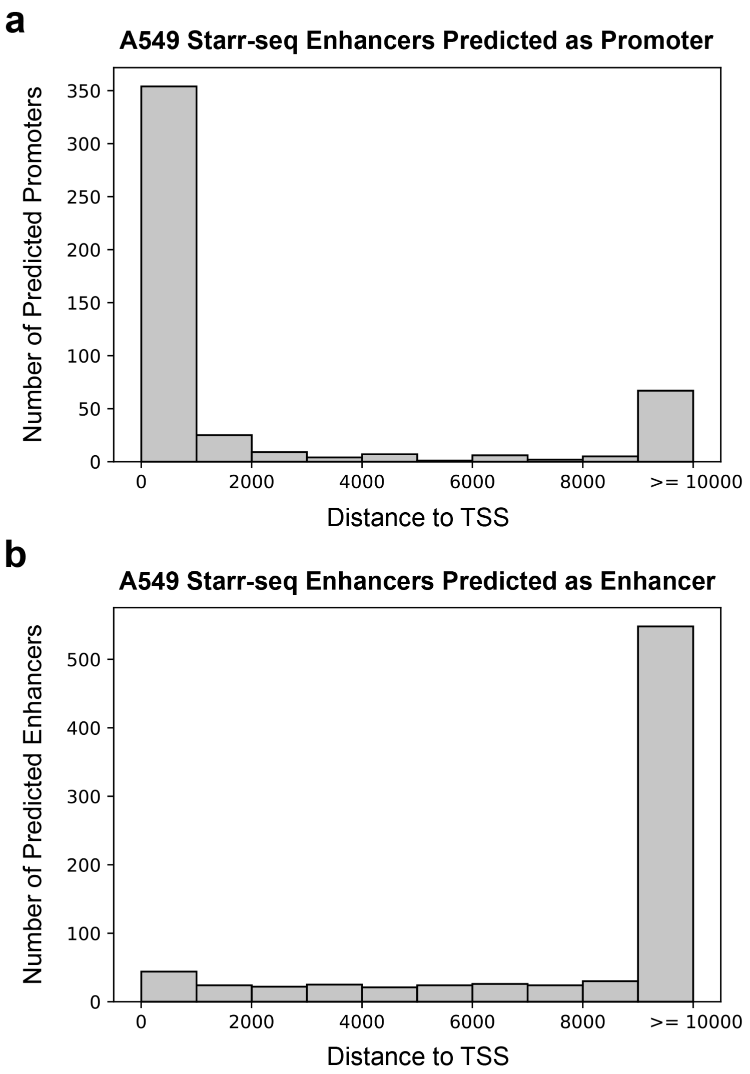


**S11 Fig. STARR-seq enhancers predicted as promoters are proximal to transcription start sites.** (**a**) Histogram of distances to the nearest TSS for STARR-seq enhancers predicted as promoters by CoRE-ATAC. Majority of predicted promoters are within 1kb of a TSS. (**b**) Histogram of distances to the nearest TSS for STARR-seq enhancers predicted as enhancers by CoRE-ATAC. Majority of predicted enhancers are distal ( >= 10kb ) from the nearest TSS. STARR-seq enhancers annotated as promoters result from the close proximity these enhancers are to a TSS.

**Supplementary Figure 12**

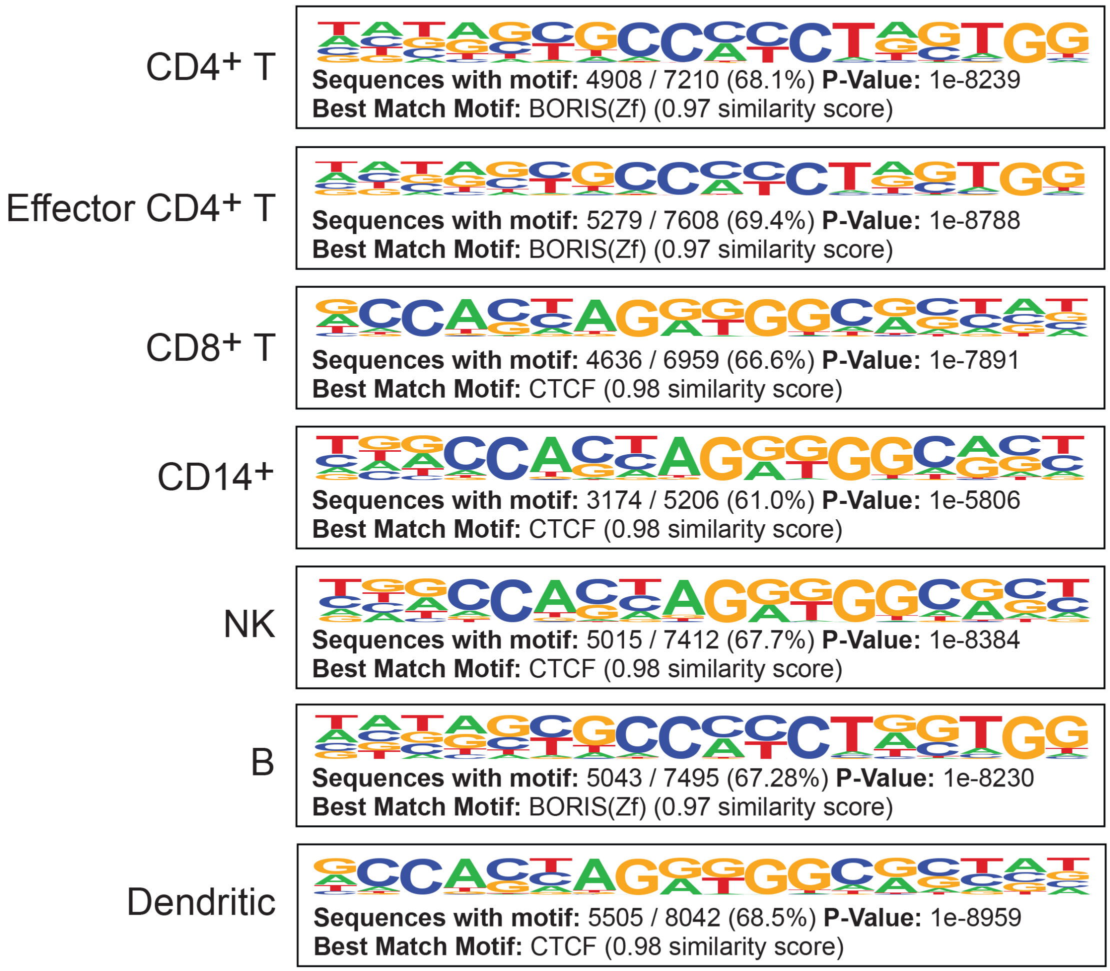


**S12 Fig. Predicted insulators in snATAC data are enriched for CTCF.** Top de novo motif enrichments for all seven cell type clusters for snATAC data. Insulator predictions for all cell types are significantly enriched for motifs with high similarity with CTCF.

**Supplementary Figure 13**

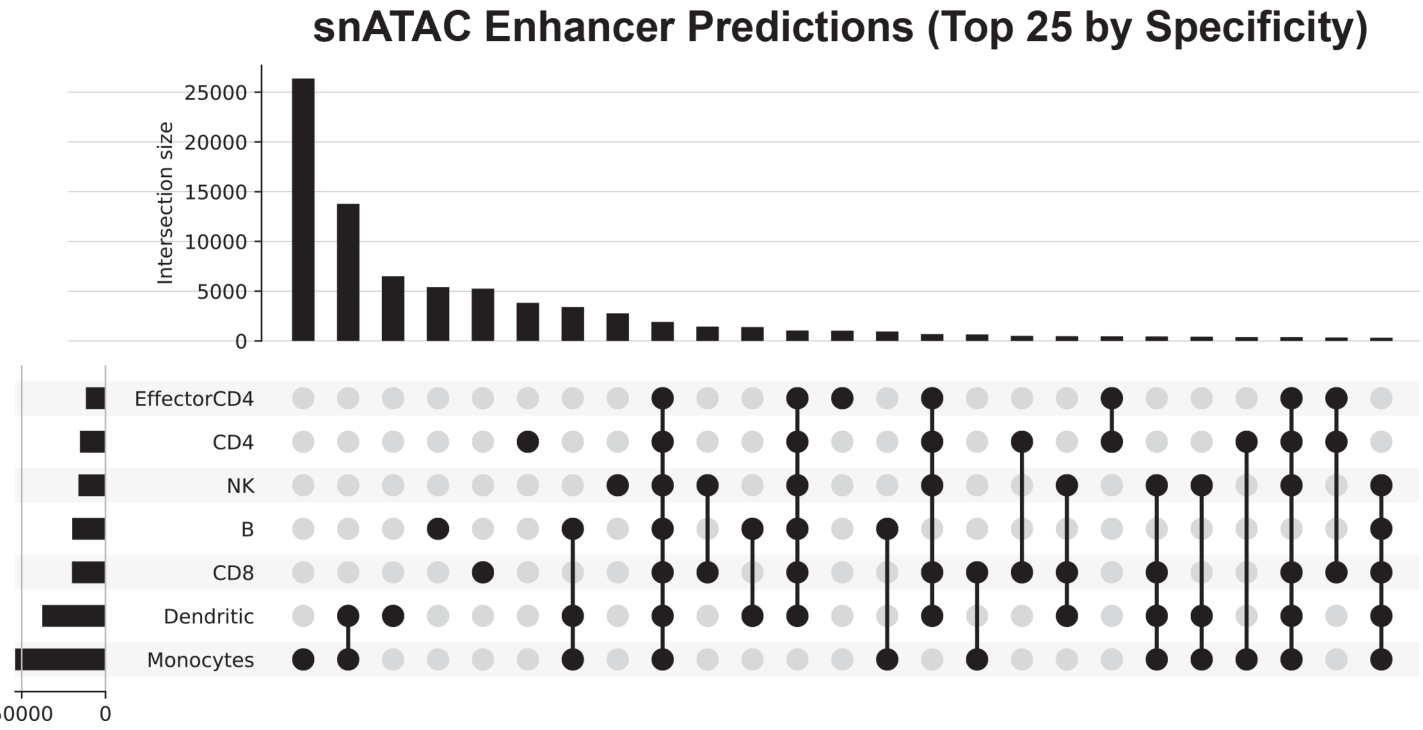


**S13 Fig. CoRE-ATAC predictions identify cell specific enhancers in snATAC-seq data.** Number of enhancers identified for the top 25 cell types and combinations by the number of enhancers predicted by CoRE-ATAC. The majority of enhancers predictions are cell-specific.

**Supplementary Figure 14**

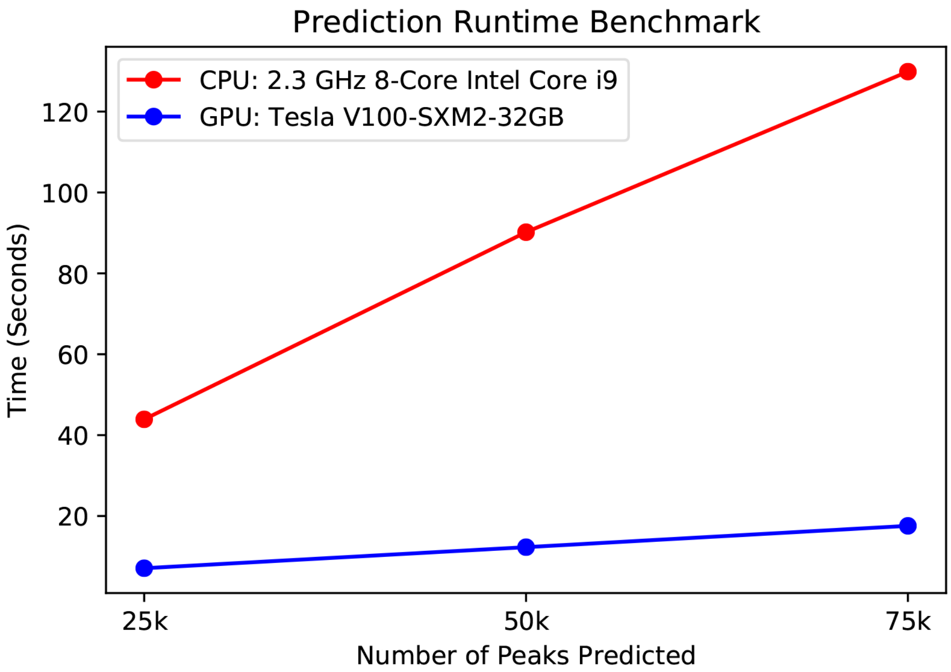


**S14 Fig. Runtime performance comparison for classifying *cis*-REs with CoRE-ATAC using CPU and GPU hardware*.*** We measured the time (in seconds) to classify cis-REs using either an intel Core i9 CPU or a Tesla V100 GPU. Comparisons revealed a linear time increase as the number of peaks increases. Predictions with the CPU take ~2 minutes for 75000 peaks, which is reasonable for users of CoRE-ATAC. GPU based method comparatively take less than 20 seconds, showing the power of using a GPU for such analyses.
